# Induction of differentiation and metabolic reprogramming in human hepatoma cells by adult human serum

**DOI:** 10.1101/180968

**Authors:** Rineke Steenbergen, Martin Oti, Rob ter Horst, Wilson Tat, Chris Neufeldt, Alexandr Belovodskiy, Tiing Tiing Chua, Woo Jung Cho, Michael Joyce, Bas E. Dutilh, D. Lorne Tyrrell

## Abstract

Tissue culture medium routinely contains fetal bovine serum (FBS). Here we show that culturing human hepatoma cells in their native, adult serum (human serum, HS) results in the restoration of key morphological and metabolic features of normal liver cells. When moved to HS, these cells show differential transcription of 22-32% of the genes, stop proliferating, and assume a hepatocyte-like morphology. Metabolic analysis shows that the Warburg-like metabolic profile, typical for FBS-cultured cells, is replaced by a diverse metabolic profile consistent with *in vivo* hepatocytes. We demonstrate the formation of large lipid and glycogen stores, increased glycogenesis, increased β-oxidation, increased ketogenesis, and decreased glycolysis. Finally, organ-specific functions are restored, including xenobiotics degradation and secretion of bile, very low density lipoprotein, and albumin. Thus, organ-specific functions are not necessarily lost in cell cultures, but might be merely suppressed in FBS. Together, we showed that cells that are representative of normal physiology can be produced from cancer cells simply by replacing FBS by HS in culture media. The effect of serum is often overseen in cell culture and we provide a detailed study in the changes that occur, provide insight in some of the serum components that may play a role in the establishment of the different phenotypes, and discuss how these finding might be beneficial to a variety of research fields.

---

Cancer cell lines are commonly used as a model to study physiological processes *in vitro*, because they are readily manipulated and can be cultured in large quantities. However, key morphological and metabolic features are often repressed or absent in rapidly dividing cells. Hepatocellular carcinoma (HCC) cell lines for example, lack key liver features, including cell polarization, VLDL secretion, and detoxification of xenobiotics^1, 2^. Additionally, cancer cells, including hepatoma cells, typically have a cancer metabolic profile^3^, using aerobic glycolysis (‘the Warburg effect’) and glutaminolyis for energy production, which is not representative of normal liver physiology and metabolism, and of the regulatory role that healthy hepatocytes play in lipid and glucose homeostasis. Although cultured primary hepatocytes are more representative of hepatocytes *in vivo* than HCC cells, they are also expensive, only available in small quantities, and difficult to manipulate (e.g. CRISPR editing). Recently we demonstrated that choosing and alternative serum source can have major implications on cell functions: when HCC cells are cultured in their native adult serum, they undergo contact inhibition and differentiate into a hepatocyte-like cell^4^. These cells can be stably maintained for months, without subculturing. We showed that in these cells key hepatic functions, like VLDL secretion are restored, and for example the production of hepatitis C virus in differentiated cells increases more than a 1000-fold, without inducing cell death, while also producing HCV particles that are more representative of the particles that are circulating in the serum of HCV infected patients^4^. In the current study, we further investigated the cellular changes that occur in HCC cells that are cultured in HS instead of FBS. We used a combination of microarray analysis, microscopic techniques, and biological assays to show that the limitations of standard HCC cultures can be overcome by changing the serum. By replacing FBS with HS in the cell culture medium, Huh7.5 cells (i) become growth arrested, obtain an epithelial, cuboid morphology and become polarized; (ii) undergo complete metabolic reprogramming, with a reversal of the cancer metabolic profile (Warburg effect and glutaminolyis); (iii) diversify other metabolic pathways, with a reduction in glycolysis, an increase in glycogen storage (glycogenesis) and higher reliance on β-oxidation; and (iv) increase mRNAs of many CypP450 enzymes and CypP450 metabolic rates and increase or restore secretory processes, like VLDL, albumin and bile secretion.

Summarizing, we show that by simply placing cells in their native adult serum, extensive reprogramming of Huh7.5 takes place, and the morphology and functions that were considered lost in cancer cell lines can be restored. We discuss the relevance of these findings for *in vitro* research, given the central role metabolism plays in various physiological processes.

## Results

### 1. Polarization, cytoskeletal organization and other morphological changes

We investigated the effect of replacing FBS by HS in tissue culture media, on cell morphology and the gene expression profile of the HCC cell line Huh7.5. We first examined overall morphological changes resulting from extended culturing in HS. HS and FBS-cultured cells where grown on transwell dishes, prepared for electron microscopy and sectioned perpendicular to the membrane surface, so that a ‘side view’ of the cell is created (Figure 1A). HS-cultured cells become cuboid, consistent with the *in vivo* hepatocyte phenotype. Moreover, their apical surface has a more pronounced epithelial character than FBS-cultured cells, with more and larger villi. HS-cultured cells are also tightly interconnected, with no open space in between, unlike their FBS-cultured counterparts. This is confirmed in higher magnification images of the cell boundaries (Figure 1B). Increased cytoplasm density and altered organelle organization were also noted in HS-cultured cells as further described in Supplemental Data 1.

**Figure 1.**
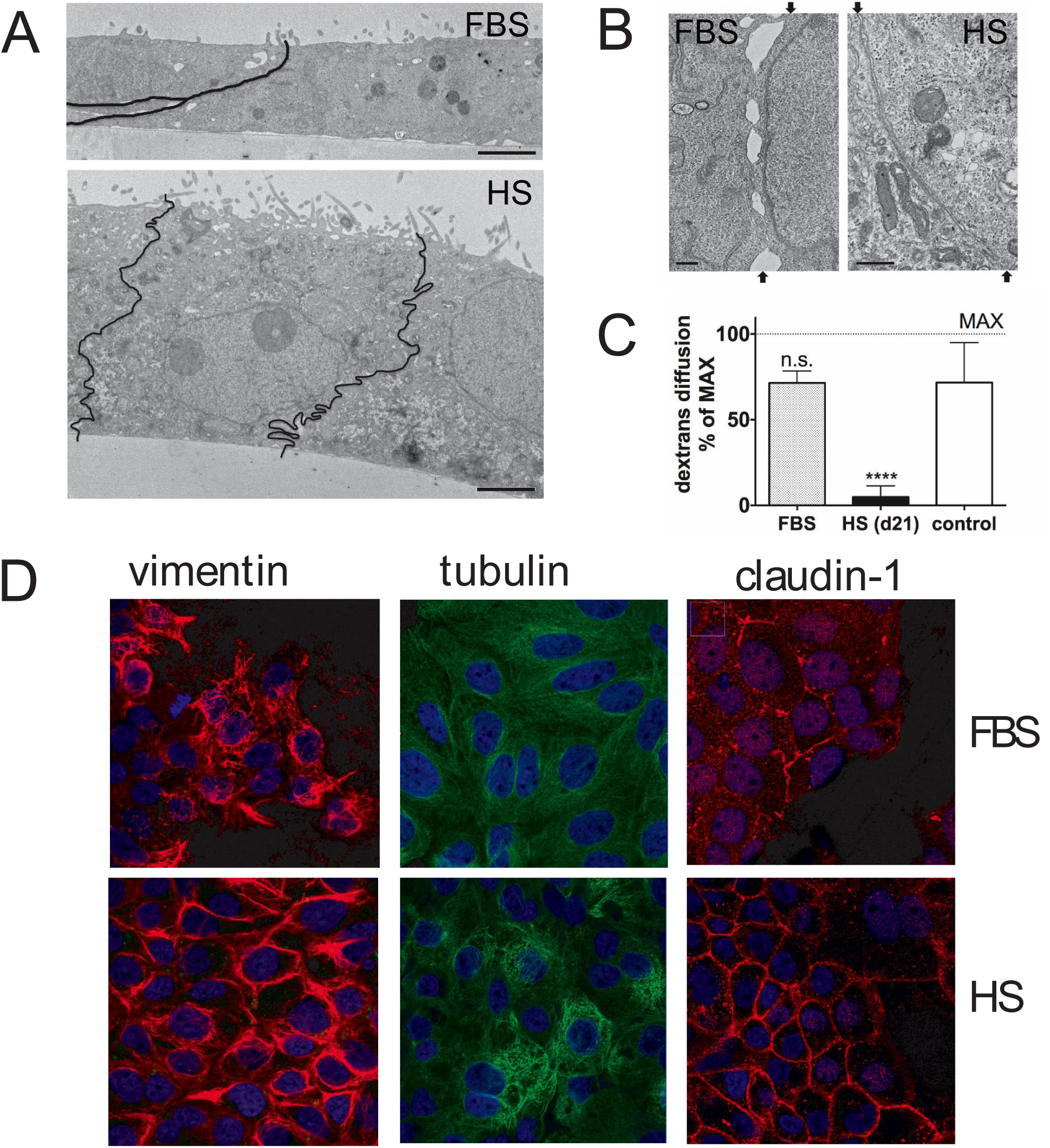
Morphology of Huh7.5 cells cultured in FBS- and HS-containing media. Electron micrographs of sagi1al sections of Huh7.5 cells that were cultured in FBS-containing media (top image) and HS-containing media (bo1om image). Black lines indicate the location of the borders between two HS cells. The images were taken at the same magnification (bar is 2 μm). Shown is a representative Figure from 2 experiments with 2 transwell dishes each. **(B)** Electron micrographs of the border between two adjacent cell in FBS (leI) and in HS (right). The arrows indicate the start and end of the border region on the image. Shown is a representative image of 3 experiments with 2 dishes each. **(C)** Dextran difiusion rate of confluent layers of FBS (FBS) and HS-cultured cells (HS d21), grown on transwell dishes. Subconfluent cultures were used as a control (control). Data are normalized to maximal difiusion rate (MAX), measured on dishes without cells. Data are presented as mean with standard deviation, from 4 independent experiments with two transwell dishes each. **(D)** Confocal imaging of FBS and HS-cultured Huh7.5 cells. Cytoskeleton components vimen2n and tubulin where stained, as well as claudin-1, a 2ght junction component. Images are representative images taken from 3 separate coverslips.

Cytoskeletal organization plays an important role in establishing polarization and cell shape. Therefore, we visualized cytoskeletal reorganization of tubulin, a microfilament, and vimentin, an intermediate filament, by confocal microscopy.

Whereas vimentin is disorganized in FBS-cultured cells, a structured organization is seen in HS-cultured cells (d21). Tubulin appears more condensed in HS-cultured cells than in FBS-cultured cells, where staining is faint and more dispersed.

We also visualized the organization of claudin-1, a major component of the tight junction complex (Figure 1D). Claudin-1 is present in FBS, but the staining is patchy, whereas in HS-cultured cells claudin-1 is distributed evenly around the entire cell, pointing at better barrier function. To test if such a barrier exists, we measured the diffusion rate of fluorescently labeled 70 kDa dextran conjugates across confluent layers of FBS-cultured cells or HS-cultured cells grown on transwell dishes. HS-cultured cells were almost impermeable to these conjugates, showing that a barrier is indeed established, whereas FBS monolayers remained permeable (Figure 1C). In conclusion, morphological hallmarks of hepatocytes, including the formation of polarized cell layers consisting of tightly interconnected cuboid cells, can be achieved in Huh7.5 cells simply by culturing them in HS instead of FBS.

### 2. Gene expression changes

Next, we used genome-wide expression arrays to investigate the overall gene expression changes of cells cultured in HS for 8, 15 and 23 days compared to cells cultured in FBS. These time points were chosen because: (i) HS-cultured cells become growth arrested after 7-10 days, (ii) around day 15 the first morphological signs of differentiation become apparent and (iii) after 21 days the differentiation process appears complete^4^. Indeed, gene expression changed significantly (p< 0.05 after multiple testing correction) in 22 to 32% of the genes (11,000-16,000 of 49,000 probes), revealing a complete cellular reprogramming upon shifting to HS media (Supplemental Data 2). Principal component analysis (PCA, Figure 2A) showed a good experimental replicability, and a clear change in expression profile upon shifting the cells from FBS to HS. To determine the similarity in gene expression between HS-cultured cells and primary hepatocytes, published hepatocyte expression profiles were projected onto the PCA, by applying the gene weights associated with PC1 and PC2 to the gene expression levels. This showed close proximity of primary hepatocytes to the later stages of the HS-cultured HCC cells, suggesting differentiation towards a hepatocyte phenotype (Figure 2).

**Figure 2.**
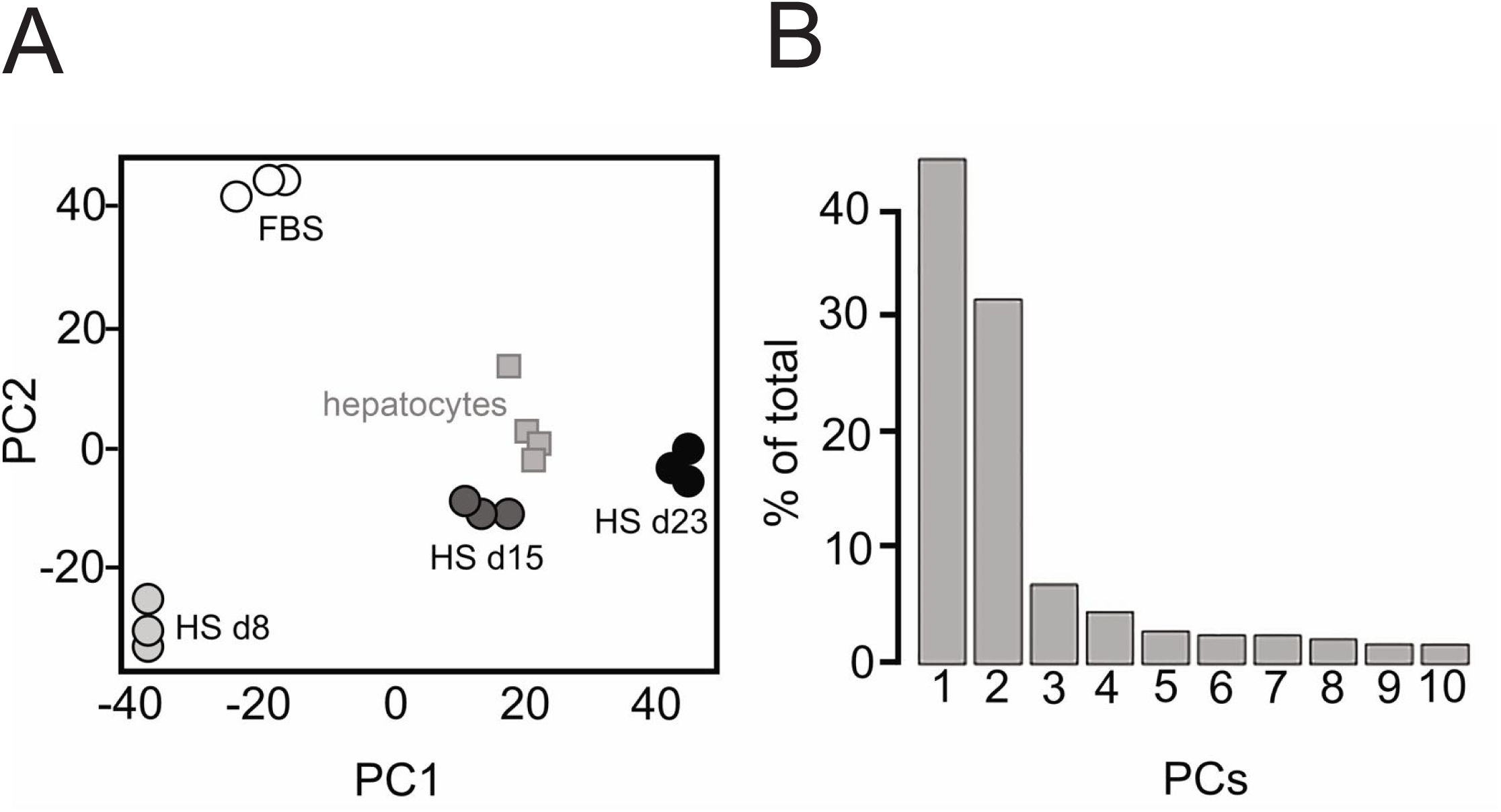
Principal component analysis. Principal component analysis (PCA) is a statistical method that is typically used as an exploratory method for data analysis, as it provides insight in the quality of a dataset, and in differences between experimental groups. **(A)** First two principal components (PCs) of triplicate genome-wide expression profiles of Huh7.5 cells in FBS and 8, 15, and 23 days aMer transfer to HS. **(B)** Scree-plot indicaing the total variance explained by the individual PCs showing that 75% of variance in gene expression is explained by the First two PCs principal components.

### 3. PAM clustering analysis

To obtain better insight into the specific genetic processes that changed as a result of culturing cells in HS, we identified clusters of co-expressed genes by applying hierarchical clustering using z-scores (Supplemental Data 3) followed by PAM (partitioning around mediods) clustering (Figure 3). Six gene expression patterns emerged, and Gene Ontology (GO) terms associated with these six patterns where determined (Figure 3, Supplemental Data 4). Changes in cell cycle and lipid metabolism gave the strongest signals related to the shift to HS. Most biological processes associated with the six clusters are consistent with our current and previous analyses^4^: cells become growth arrested (cluster 1, 4, 5) and lipid droplets, as well as VLDL secretion were increased (cluster 1, 6). We also observed a de-repression of apoptosis (cluster 2). Cancer cells often repress apoptosis, and the observed changes may reflect the loss of their proliferative character.

**Figure 3.**
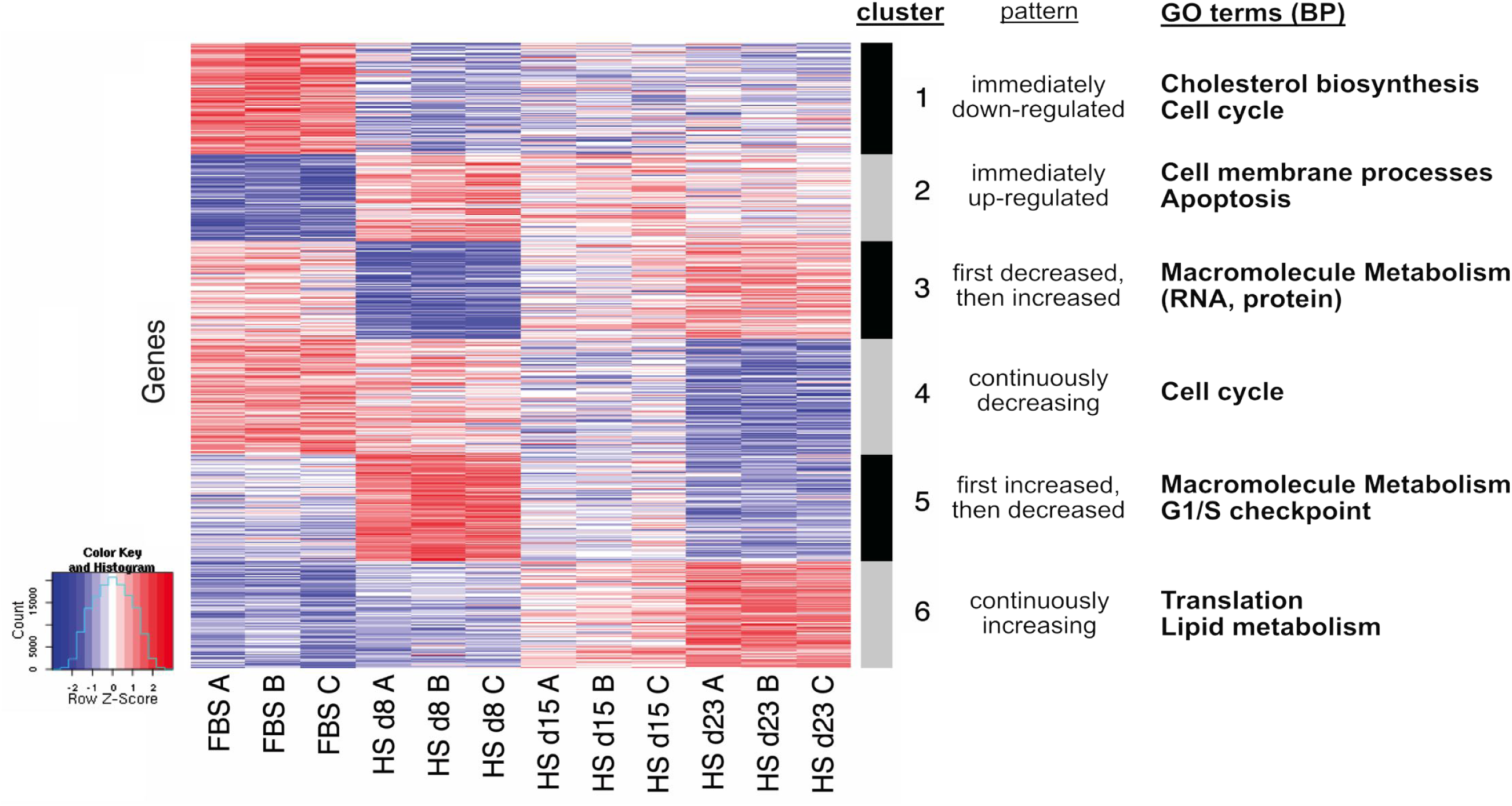
PAM clustering. PAM clustering (partioning around medoids) of the microarray data into 6 clusters. Samples are on the x-axis while genes are on the y-axis. Gene expression values were standardized across the samples into Z-scores. Red indicates higher expression relative to other samples, blue indicates lower expression. Six paHerns of expression emerged from this analysis, and the Gene Ontologies terms/ Biological Processes (GO terms/ BP) associated with these clusters were determined (Supplemental Data 4) and summarized on the right of the Figure.

Thus, we further examined the metabolic changes occurring in HS-cultured cells that are predicted to accompany the transition from a proliferating to a differentiated and growth-arrested state of the cell.

### 4. Metabolic reprogramming: reversal of the Warburg effect and metabolic diversification

Proliferating cells often display a ‘cancer metabolism’ profile, first described by Otto Warburg in 1924, which includes reduced levels of oxidative phosphorylation and mitochondrial activity, higher dependence on aerobic glycolysis and glutaminolysis for ATP production, and increased generation of biosynthetic intermediates that are essential for the production of macromolecules (phospholipids, nucleotides, proteins) to support cell proliferation (reviewed in^3, 5-7^). The metabolic reprogramming that occurs during the Warburg effect is tightly regulated. Key regulators include pyruvate dehydrogenase kinase 1 (PDK1), the lactate dehydrogenase A/B ratio (LDHa/LDHb), and monocarboxylic acid transporter 4 (MCT4), as further explained in Supplemental Data 5.

To test our hypothesis that cancer metabolism is reversed in HS-cultured Huh7.5 cells we compared HS- to FBS-cultured cells using a combination of gene expression analyses, measurement of end-point metabolites, electron microscopy and biological assays. The first indication that HS-cultured cells rely less on aerobic glycolysis than their FBS-cultured counterparts, is the much slower acidification of the cell culture media, as indicated by medium colour. In line with this, and with the reversal of the Warburg effect, mRNA of LDHa was decreased and LDHb was increased, shifting the reaction away from lactate production (Figure 4A). Additionally, mRNA of lactate transporter MCT4 was also decreased in HS-cultured cells, as well as strong downregulation of PDK1, which regulates pyruvate uptake by mitochondria (Figure 4A, Supplemental Data 6)

**Figure 4.**
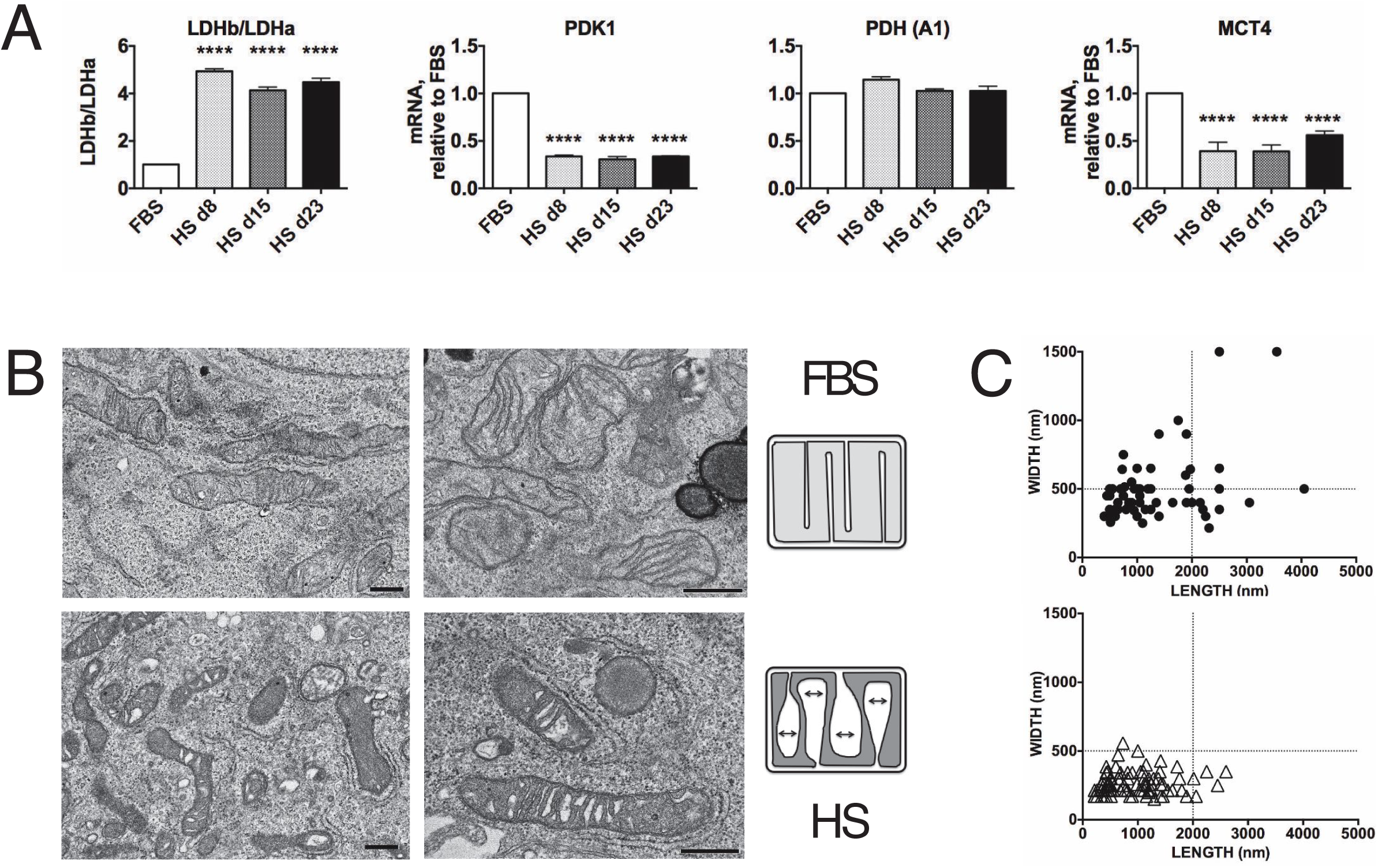
Reversal of the Warburg effect. Relative expression of genes involved in the regulation of the Warburg effect. Data were obtained from mRNA isolated a time series with 3 biological replicates each, and at least 2 measurements per biological replicate. Data are depicted as a fold change over FBS. Depicted is mean with s.d., statistical significance was determined by ANOVA/ DunneI (one-sided, with multiple comparisons correction). P values ranges are depicted as asterisks: *: P< 0.05, **: P< 0.01, ***: P< 0.001, ****: P≤0.0001, n.s: P> 0.05 **(B)** Mitochondrial morphology in cells that where cultured in FBS (top images) and cells that were cultured in HS-containing media (bottom images), and schematic representation of mitochondrial morphology. Shown is a representative image, morphology was consistent in 3 separate EM experiments with 2 cell culture dishes each. **(C)** Measurement of length versus width of mitochondria on electron micrographs. Top panel: FBS, boIom panel: HS. Although these measurements do not represent the real size for the mitochondria, a similar size distribution is expected when mitochondria are the same size, when a high amount of mitochondria is analyzed. In this analysis, mitochondria in HS-cultured cells are narrower than those in FBS-cultured cells. Depicted are individual measurement of mitochondria on electron micrographs, taken in 3 separate experiments with 2 cell culture dishes per experiment.

Changes in mitochondrial morphology also might suggest increased rates of oxidative phosphorylation in HS-cultured cells. In FBS-cultured cells, the mitochondrial matrix appeared lighter than in HS, where it was condensed and dark with large spaces between the cristae (Figure 4B). Hackenbrock^8-10^ described similar mitochondrial transitions in isolated mitochondria, and linked them to electron transport chain activity: condensed mitochondria are indicative of high activity, whereas orthodox mitochondria indicate low activity. Other studies have since confirmed the link between mitochondrial activity and mitochondrial morphology^11-13^. High oxidative phosphorylation rates are also associated with mitochondrial narrowing^13^, and we made similar observations in HS-cultured cells (Figure 4C). Although the physiological role of the mitochondrial heterogeneity remains largely unclear, matrix density may influence diffusion rates of metabolites in and out of the mitochondria^14-16^, for example of ATP and ADP.

We investigated which metabolic pathways changed as Huh7.5 cells shifted away from their cancer metabolism. Figure 5A shows that fermentation and glutaminolysis were decreased in HS-cultured cells, consistent with the reversal of cancer metabolism. Additionally, (i) all but one glycolysis enzymes were down-regulated, including the rate-limiting step. The one enzyme that is up-regulated is also involved in gluconeogenesis, and the mRNAs of enzymes in the gluconeogenesis pathway were generally up-regulated (data not shown). (ii) Most glycogenesis enzymes, including the rate-limiting step were up-regulated, indicating that HS-cultured cells convert large amounts of glucose to glycogen, relying less on glucose for ATP production. This is supported by the presence of large glycogen deposits within the cell, as is shown in Figure 5B, bottom image (marked G). Glucose use was not altered in HS-cultured cell compared to FBS-cultured cells (Figure 5D, left panel). (iii) In HS-cultured cells lipid droplets size was increased (Figure 5B, top two panels^4, 17^). β-oxidation is in part regulated by the availability of lipid stores and metabolic analysis supports an increase in β-oxidation rate: many enzymes involved in TG (Triacylglycerol) degradation and β-oxidation are increased, including the rate regulating step of β-oxidation, CPT-1 (carnitine-palmitoyl transferase 1). Increased protein levels of CPT-1 (approximately 7-fold) were confirmed by western blot (Figure 5C). (iv) Acetyl CoA produced by β-oxidation is partially converted to ketone bodies in the liver of healthy individuals, which play a critical role in normal energy homeostasis^18^. Formation of ketone bodies is therefore often used to estimate the rate of β-oxidation. Ketogenesis was also increased in HS-cultured Huh7.5 cells (Figure 5A), which was confirmed by NMR analysis of metabolic end-products in HS-cultured cells. Acetoacetate and 3-hydroxybuyrate, the two main ketone bodies produced during ketogenesis, where significantly increased in HS cultured cells (Figure 5D). This further supports increased levels of β-oxidation in HS serum cultured cells.

**Figure 5.**
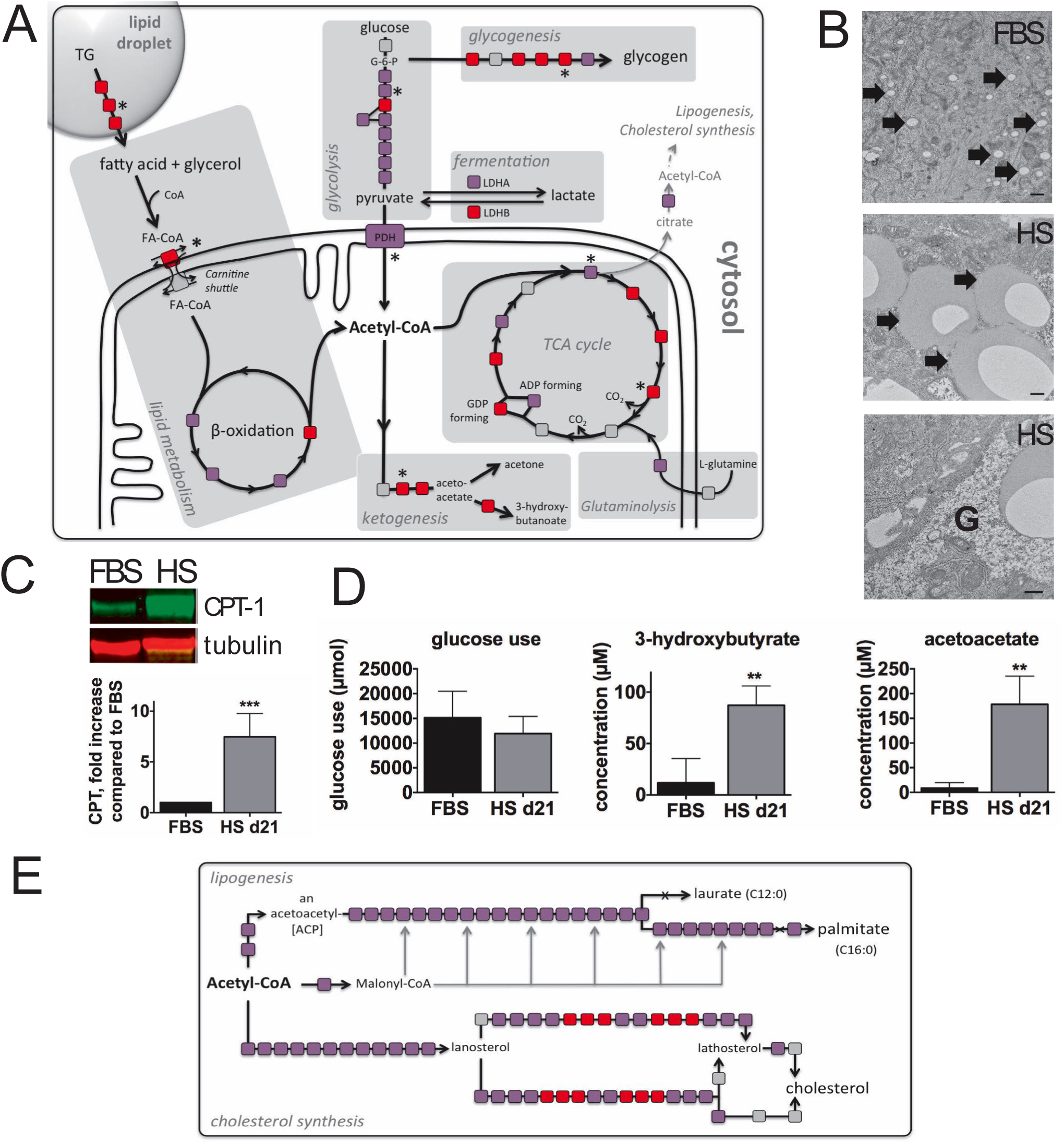
Metabolic mapping. Analysis of several metabolic pathways using HumanCyc. Each enzyme in a biosynthe;c pathway is depicted as a square. Changes in transcription levels are colour-coded: red indicates an activated gene (P< 0.05), purple indicates a deactivated gene (P< 0.05) and grey indicates no change. Reactions marked with “X” indicates no microarray data were available. Asterisks indicate the (predicted) rate limiting or rate regulating steps of the metabolic pathway. **(B)** Lipid droplets (indicated by arrows) in FBS and HS-cultured cells and presence of glycogen stores in HS-cultured cells (indicated by G). Shown are represantive images taken in 3 separate experiments with 2 cell culture dishes each. **(C)** Licor blot of CPT-1 (carnitine palmitoyl transferase-1) and tubulin. Shown is a representative image of 5 blots, quantitation of those 5 blots is represented in the boVom panel. Values are depicted as fold increase over FBS (n=5, mean ± s.d.). Statistcal significance was calculated using a t-test. ***: P< 0.001. **(D)** Quantitation of ketone bodies present in the cell culture supernatant of HS and FBS-cultured cells, by NMR. n=4, depicted is mean± s.d.. Statistical significance was calculated using a t-test. **: P≤0.01. (E) Enzymes involved in palmitate and cholesterol synthesis and their differential expression paVerns. See (A) for further details.

Finally, consistent with the reversal of a proliferative metabolic profile, mRNAs of pathways that produce ‘building blocks’ for cell growth and proliferation, specifically cholesterol synthesis and lipogenesis, are decreased (Figure 5E). In these pathways, citrate that is produced in the mitochondria is exported to the cytosol, where it is converted to Acetyl-CoA. Acetyl-CoA can then be incorporated into cholesterol and fatty acids, as depicted in Figure 5E. mRNAs of most genes in these pathways are decreased in HS-cultured cells. Malonyl-CoA is also produced in the lipogenesis pathway and inhibits CPT-1, and thereby β-oxidation. Thus, its decreased production in HS-cultured cells is in line with the activation of β-oxidation in HS-cultured cells, and metabolic reprogramming in general.

Summarizing, our analyses show that the metabolism in HS-cultured cells shifts away from cancer metabolism (glutaminolysis and the Warburg effect) and glycolysis, in favor of glycogen storage. Lipid stores are increased, as are β-oxidation and ketogenesis. The increase in lipid droplet size is probably the result of the activation of TG biosynthesis and storage pathways, that are regulated by LXRα and PPARγ, which in turn are both upregulated in HS-cultured cells^4^. The presence of native lipoproteins in the human serum may also play a role: when we used lipoprotein free human serum, differentiation proceeded as expected, but the increase in lipid droplet size was largely prevented (not shown).

### 5. Xenobiotics biodegradation and metabolism

As cells move away from the state of proliferation and cancer metabolism, surplus nutrients become available for storage, and to reinstate secretory processes like VLDL secretion. Indeed, we showed previously that VLDL secretion is completely absent in FBS-cultured cells, but as cells become growth arrested and differentiate, VLDL secretion is gradually restored^4^. Here, Kyoto Encyclopedia of Genes and Genomes (KEGG) pathway enrichment analysis supports the re-establishment of lipid and carbohydrate metabolism as described in the previous sections, as well as re-establishment of secretory processes (Table 1, Supplemental Data 7). Enriched pathways included ‘steroid hormone synthesis’ (KEGG pathway hsa00140), ‘degradation of xenobiotics by cytochrome P450’ (KEGG pathway hsa00980) and ‘ascorbate and aldarate metabolism’ (KEGG pathway hsa00053). Steroid hormones derive from cholesterol and are secreted by various organs including the liver. Ascorbate and aldarate metabolism is central to many conversions of glucose, including nucleoside synthesis and pentose interconversions, which itself was significantly increased after 15 days in HS media. The most notable changes involved ‘Xenobiotics biodegradation and metabolism’. Three pathways in this cluster were significantly increased (Table 1) and several detoxification enzymes were present in the top-25 transcripts with increased expression. Notably, the enzyme with highest transcriptional increase was sulfotransferase 1E1 which catalyzes the sulfate conjugation of xenobiotics, facilitating removal (Supplemental Data 6).

**Table 1:** Significantly increased KEGG pathways (p< 0.05)

|  | **<u>KEGG ID</u>** | **<u>Pathway name</u>** | **<u>Fold</u>**<br>**<u>change</u>** | **<u>p-value</u>** |
| --- | --- | --- | --- | --- |
| HS D8 | hsa00980 | Metabolism of xenobiotics by cytochrome P450 | 3.65 | 0.0028 |
|  | hsa00140 | Steroid hormone biosynthesis | 3.90 | 0.0040 |
|  | hsa00053 | Ascorbate and aldarate metabolism | 4.27 | 0.0317 |
| HS D15 | hsa00140 | Steroid hormone biosynthesis | 8.18 | 6.16E-008 |
|  | hsa00980 | Metabolism of xenobiotics by cytochrome P450 | 6.56 | 6.45E-007 |
|  | hsa00982 | Drug metabolism - cytochrome P450 | 5.22 | 4.69E-005 |
|  | hsa00830 | Retinol metabolism | 5.30 | 0.0001 |
|  | hsa00983 | Drug metabolism - other enzymes | 4.89 | 0.0012 |
|  | hsa00053 | Ascorbate and aldarate metabolism | 6.52 | 0.0029 |
|  | hsa00040 | Pentose and glucuronate interconversions | 5.30 | 0.0063 |
|  | hsa00500 | Starch and sucrose metabolism | 3.92 | 0.0084 |
|  | hsa00860 | Porphyrin and chlorophyll metabolism | 3.94 | 0.0179 |
|  | hsa00514 | Other types of O-glycan biosynthesis | 3.68 | 0.0224 |
| HS D23 | hsa00980 | Metabolism of xenobiotics by cytochrome P450 | 5.18 | 8.63E-007 |
|  | hsa00140 | Steroid hormone biosynthesis | 5.46 | 3.60E-006 |
|  | hsa00982 | Drug metabolism - cytochrome P450 | 4.26 | 4.22E-005 |
|  | hsa00830 | Retinol metabolism | 4.42 | 6.90E-005 |
|  | hsa00053 | Ascorbate and aldarate metabolism | 5.44 | 0.0018 |
|  | hsa00983 | Drug metabolism - other enzymes | 3.81 | 0.0021 |
|  | hsa00040 | Pentose and glucuronate interconversions | 4.42 | 0.0048 |
|  | hsa00500 | Starch and sucrose metabolism | 3.14 | 0.0119 |
|  | hsa00860 | Porphyrin and chlorophyll metabolism | 3.29 | 0.0168 |
|  | hsa00514 | Other types of O-glycan biosynthesis | 3.08 | 0.0220 |

To confirm the increase in xenobiotics metabolic capacity, we focused on a panel of Cytochrome P450 (CypP450) enzymes, the key enzymes that degrade xenobiotics (Figure 6A, Supplemental Data 8). All CypP450 enzymes had increased mRNA levels in HS (1.5-60 fold), with the exception of Cyp4A11. Cyp4A11 is involved in the detoxification of lipid products, when β-oxidation is absent or decreased. Thus, this finding is consistent with the observed increase in β-oxidation in HS-cultured cells. Transcriptional levels of other proteins involved in degradation and removal of xenobiotics were also determined, and findings were generally consistent with increased degradation and removal of toxic compounds (Supplemental Data 9).

**Figure 6.**
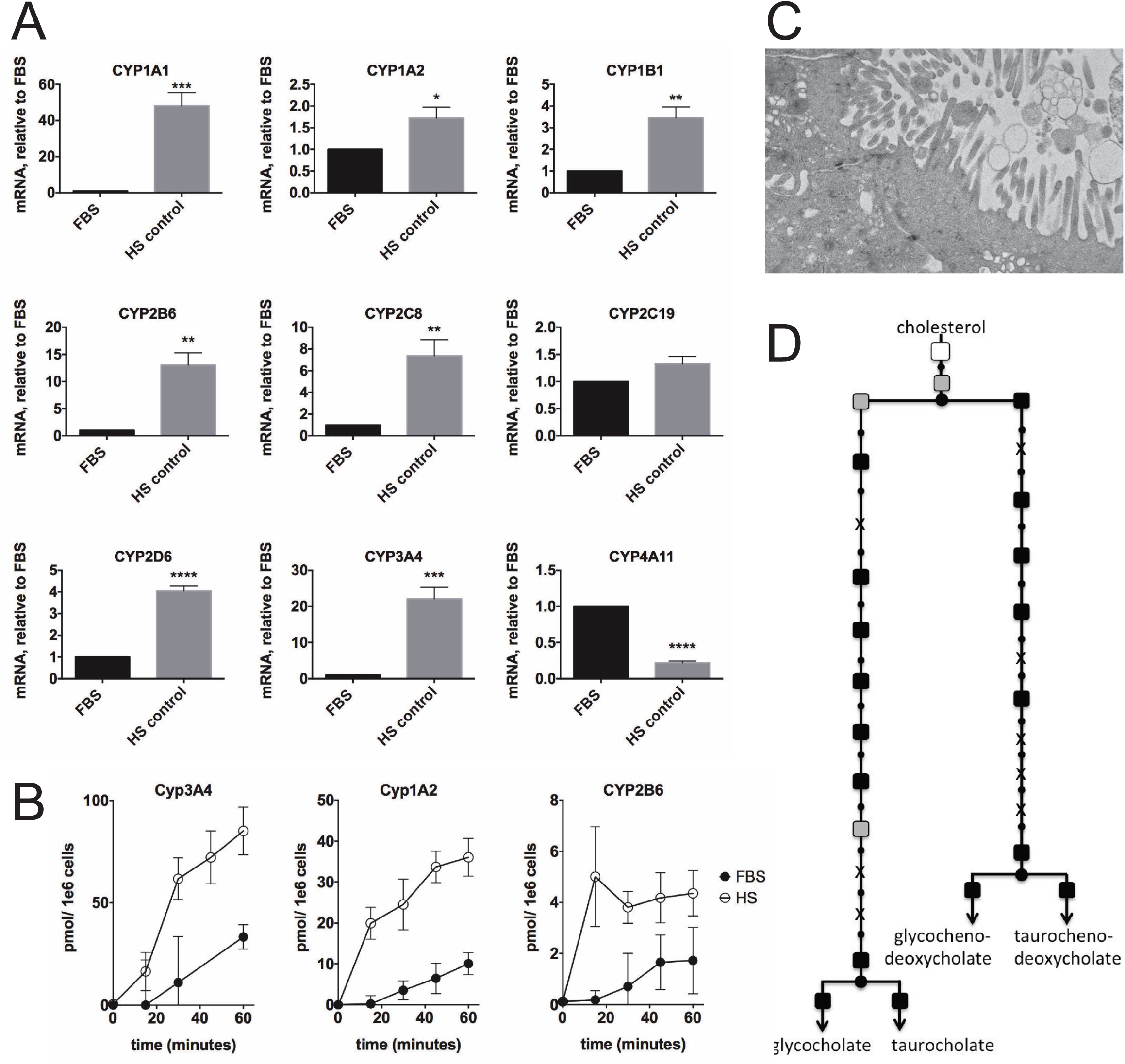
Cytochrome P450 metabolism and bile production. mRNA of different cytochrome P450 (CYP) genes, as determined by quantitative PCR. Data are depicted as mean±s.d. n=4 (4 biological replicates with duplicate measurements each). Statistical significance was calculated using Student’s t-test. P values ranges are depicted as asterisks: *: P< 0.05, **: P< 0.01, ***: P< 0.001, ****: P≤0.0001 **(B)** Activity of selected CYPs, measured with a substrate cocktail assay. Shown is a representative time-series, of 3 independent experiments with duplicate cell culture dishes each. **(C)** Electron micrograph of a structure resembling a bile canalicular surface. Bar represents 500 nm. Shown is a representative Figure from 2 experiments with 2 transwell dishes each. **(D)** Metabolic mapping of microarray data, of enzymes involved in bile acid synthesis. Changes in transcription levels are colour-coded: black indicates an activated gene (P< 0.05), white indicates a deactivated gene (P< 0.05) and grey indicates no change. Reactions marked with “X” indicates no microarray data were available.

To functionally confirm the changes observed at a transcriptional level, we examined the rates of xenobiotics degradation in FBS- and HS-cultured cells (Figure 6B). The intrinsic, non-induced rates of degradation by Cyp3A4 (testosterone), Cyp1A2 (methoxy-resorufin) and Cyp2B6 (buproprion) were increased approximately 4-, 6- and 6-fold respectively.

Many products resulting from CypP450 activity are removed from the liver in the bile, through the bile canalicular surface, which is only established in polarized cells. Bile synthesis is also dependent on the availability of excess cholesterol. Figure 6C shows the formation of structures that have a high resemblance to bile canaliculi in HS-cultured cells. Most mRNAs of enzymes involved in bile synthesis are increased (Figure 6D), indicating higher levels of bile acid secretion. Similar observations were reported elsewhere in HepG2 cells which were cultured in human serum^19^.

## Discussion

When tissue and cell culture was initially developed in the early to mid 1950s, the lack of cell growth was a major problem, which was overcome by the addition of fetal animal sera or embryo extracts. The first human cell line for example, HeLa cells, was grown in chicken plasma clot, mixed with saline, human umbilical cord serum and bovine embryo extract^20^. Eventually, fetal bovine serum (FBS) became the serum of choice, because of its abundant availability and excellent growth promoting capacities. Undeniably, great progress has been made since cell cultures were first developed, however, it has also become increasingly clear that rapidly dividing cells are not representative of normal functional cells in an organ or organism, including the morphology of the cell, metabolism and organ-specific functionality.

In this study we used a human hepatocellular carcinoma cell line, Huh7.5, and showed that culturing these cells in their native adult serum, instead of FBS, results in growth arrest and drastic changes in morphology and intracellular organization, metabolic reprogramming and re-establishment of organ-specific functions, like VLDL secretion, albumin secretion and detoxification of xenobiotics. Significant changes in transcription in 22-32% of the genes are indicative of extensive cellular reprogramming. We showed that metabolism shifts from a cancer-like profile, to a profile that is more representative of hepatic metabolism, that includes glycogenesis, higher β-oxidation rates, lower glycolysis rates, as well as restoration of processes like VLDL secretion^4^, degradation of xenobiotics by cytochrome P450s, and bile secretion (this study,^19^). This refutes the notion that cell lines derived from cancers have lost their ability to undergo contact inhibition and much of their organ-specific functionality. We showed in Huh7.5 cells, that most of these lost functions, and a normal metabolic profile, are merely suppressed in the presence of a growth inducing serum, like FBS, and can be restored by culturing cells in their native adult serum. Other cell lines appear to undergo similar changes when cultured in their native adult serum, although this is not a universal effect. For example, other hepatocellular carcinoma cell lines (HepG2, Huh-7) undergo similar transitions (unpublished observations, ^19^). HeLa cells in our hands did not tolerate culturing in human serum, while A549 cells, a adenocarcinomic human alveolar basal epithelial cell line, did thrive in HS, stopped dividing and underwent drastic morphological changes.

It is highly unlikely that one or only a few factors in HS are responsible for transcriptional changes in 32% of the genes of the cells. Like FBS, HS is a complex mixture of components and undoubtedly many serum components play a role in the establishment of the different phenotypes in different sera. We have, in this and in previous studies determined some of the groups of components in FBS or HS that contribute to the different phenotypes. So far we have found that (i) the lack of growth factors in adult serum relative to fetal serum induces growth arrest, (ii) the enrichment of certain differentiation inducting factors in HS relative to FBS helps with differentiation, and (iii) the specific lipid composition of human relative to bovine sera may play a role. Cell proliferation slows down immediately after Huh7.5 cells are transferred to HS, and after 7-10 days proliferation has completely stopped. Tight junctions are dissociated during cell proliferation so the absence of growth factors enables formation of cell/cell contacts and tight junctions, resulting in proper cell morphology. Growth factors also regulate oncogenes, which in turn influence metabolism, e.g. oncogenes c-Myc and p53 directly regulate mitochondrial respiration^21^,^22^, and are amongst the most common alterations in human cancers, including in Huh7.5 cells^23, 24^. c-Myc increases LDHa, whereas p53 controls the rate limiting step of glycolysis^21^. Thus, the lack of oncogene stimulation, through the reduced levels of growth factors in adult serum (like HS) might play a direct role in the observed metabolic reprogramming.

The native character of HS may also be important, as other adult sera do not result in the same changes: Huh7.5 cells cultured in *adult* bovine serum (ABS) undergo growth arrest and up-regulate tight junction and cell-cell contact components^4^, but otherwise lack hepatocyte functionality like albumin and VLDL secretion. Conversely, human umbilical cord blood serum (CBS), the closest human equivalent of FBS, induces high cell proliferation rates, but also increases Huh7.5 functionality as shown by increased levels of albumin secretion. However, VLDL secretion is not restored in CBS-cultured cells^25^. Likely, VLDL secretion co-depends on the presence of sufficiently large TG stores (lipid droplets), and may thus also depend on a non-proliferative metabolic profile/ growth arrest.

Growth factors that are remarkably higher in human serum (CBS and HS), compared to FBS, are IGF-1 (insulin like growth factor 1) and several IGFBPs (IGF binding proteins)^26^. IGF-1 is increased approximately 20 and 40-fold in HS and hUCBS respectively, relative to FBS. IGF-1 and IGFBPs both play a role in hepatocyte differentiation, with IGFBPs modulating and prolonging the activity of IGF-1. Consistent with their role in differentiation, IGF-1 and IGFBPs impede the aggressive growth of certain liver cancers^27-29^. IGFBP-3 is one of the factors listed in the top-25 genes with the greatest increase in transcription in our microarray analysis (Supplemental Data 6).

In addition to differences in growth factor level and composition, the composition of the lipids in HS is also markedly different from FBS. Lipoproteins in bovine serum mainly contain saturated fatty acids, whereas lipoproteins in human serum are enriched in lipids containing unsaturated fatty acids, particularly oleic acid (18:1)^30^. The presence of oleic acid is significant, as oleic acid is often added to cells to stimulate lipoprotein secretion^31^, which cannot be achieved by addition of most other fatty acids.

These findings are relevant for many fields. In general HS-based cell culturing may increase the usefulness of human cancer cell lines by establishing affordable, scalable and physiologically relevant cell culture models. In the case of Huh7.5 cells, they enable the study of normal hepatocyte physiology, but also the transition from normal physiology to the diseased state, for example as a result of chronic viral infections like HCV infection. Metabolic (dys)regulation plays a central role in cancer, immune regulation, metabolic syndrome, diabetes and fatty liver disease, and availability of cells with a ‘normal’ metabolic profile is advantageous in deciphering the factors that lead to the diseased state. But foremost, from a cell biology perspective, the profound changes that we observed, simply as a result of changing serum in cell cultures, where unexpected and surprising, and shows that much remains to be learned about the effect of native sera *and* of FBS.

## Acknowledgements

The authors thank Dr. C. Rice (Rockefeller University) for providing us with the Huh7.5 cell line.

Funding: BED was supported by the Netherlands Organization for Scientific Research (NWO) Vidi grant 864.14.004.

## Methods

### Standard Cell culture conditions for proliferation

Huh7.5 cells were a kind gift of Dr. C. Rice, Rockefeller University, New York, USA), and were maintained according to the protocols provided. In short, cells were maintained in DMEM/10% FBS /penicillin/ streptomycin. The cell line used in this study has a low passage number, cell were never replated at a ratio higher than 1:4, and after approximately 20-30 passages, cells were discarded, and a new vial was thawed.

### Maintenance of cells in human serum

Since the use of human serum results in growth arrest^4^, cell cultures were normally maintained in FBS-containing media as described above. At the time of transfer to human serum cells were trypsinized, trypsin was inactivated with DMEM/10% FBS/ penicillin/ streptomycin and cells were centrifuged at 300g. Cell pellets were then resuspended in DMEM/2% HS/ penicillin/ streptomycin, and plated at a density of 30-50%. At confluency cells were trypsinized once more, plated at a density of 50%, and then left to form confluent layers of undividing cells. Although cells can be trypsinized/ sub-cultured for approximately 7-10 days, continued sub-culturing leads to cell death, as cells appear to loose their ability to attach to untreated cell culture plastic.

### Microarray analysis

Cells were cultured (3 biological replicates/ flasks) either in FBS, or in human serum for 8, 15 or 23 days. These days were chosen because HS-cultured cells undergo growth arrest around day 7-10 post transfer, the first morphological signs of differentiation appear around day 14, and the differentiation process appears complete after 21 days in human serum containing media. Cell lysates were then prepared for microarray analysis of Affymetrix GeneChip® PrimeView^(tm)^ Human Gene expression Array cartridges according to the instructions of the manufacturer. Microarray data were deposited in the GEO repository with Accession Number GSE87684 (GEO)

All expression data analyses were performed using the R statistical program (R Development Core Team; R: A Language and Environment for Statistical Computing (R Foundation for Statistical Computing);^32^). Probe intensities were normalized with the “affy” R package from the Bioconductor project^33^, using the “Robust Multiarray Averaging” (RMA) algorithm^34^. Differential expression analysis was performed using the “limma” R package^35^, which fits a linear model to the expression data. Principal Components Analysis was performed using the R “prcomp” function. For the clustering analysis gene expression levels were converted to z-scores, standardized across the samples, to eliminate the differences in absolute gene expression levels, and focus on the pattern of changes. An initial hierarchical clustering was performed using Euclidean distances and average linkage in order to get an idea of how many clusters were present in the data. Based on this the expression data were clustered into 6 clusters using the Partitioning Around Medoids (PAM) algorithm, which requires a predetermined number of clusters to be specified. Gene Ontology term enrichment calculations were performed using the “goseq” R package^36^.

### Metabolic modeling

For our metabolic modeling purposes we use a recently constructed stoichiometric metabolic network for human, called Recon2^37^. The boundaries of exchange reactions were left at default values and dead-end reactions (i.e. the reactions whose product metabolites are not used for any other reaction or for growth) were removed. We integrate the measured gene expression values with this model using a program called Metabolic Adjustment by Differential Expression (MADE)^38^. MADE uses the p-values and fold changes obtained in a differential gene expression analysis to identify enzymes with significantly different expression between conditions, and looks for the best fit with the metabolic network. The program predicts for each gene if it is activated, deactivated or stable between conditions and time points, such that each prediction is consistent with the network structure. This can be visualized as activation and inactivation of the corresponding reactions using the online tool HumanCyc^39^

### Transmission Electron microscopy (TEM)

For conventional transmission electron microscopic study, cells were either cultured in FBS and grown to confluence, or cultured in HS-containing media, grown to confluence and allowed to differentiate. Some FBS and HS-cultured cells were grown on a membrane of BD Falcon cell culture inserts (cat. #353180, BD Biosciences, Canada), in order to obtain perpendicular sections of cells on the membrane surface. 2x conventional TEM fixative (mixture of 4% glutaraldehyde, 2% paraformaldehyde, 0.2 M sucrose and 4 mM CaCl_2_ in 0.16 M sodium cacodylate buffer, pH 7.4) was added to the cell media to make optimally diluted fixative. Pre-fixation was performed at 37 °C for 1 hour. Following fixation, cells were washed with 0.05 M sodium cacodylate buffer (cat. #11654, Electron Microscopy Sciences, USA) to remove residual aldehyde from cells. For lipid fixation, cells were post-fixed with 1% ice-cold osmium tetroxide (OsO_4,_ cat. #19140, Electron Microscopy Sciences, USA) in 0.05 M sodium cacodylate buffer. Following lipid fixation, cells were again washed with 0.05 M sodium cacodylate buffer. To increase contrast of cell membrane and subcellular membrane (en bloc stain), cells were treated with 1% uranyl acetate (cat. #22400-4, Electron Microscopy Sciences, USA) in 0.1 M sodium acetate buffer (pH 5.2) for 15 min. Cells were washed with 0.1 M sodium acetate buffer followed by Milli-Q filtered water and then dehydrated with ascending ethanol series (30%, 50%, 70%, 80%, 90%, 95% and 100%). Cells on membranes were gradually infiltrated with Spurr’s resin (cat. #14300, Electron Microscopy Sciences, USA). Several pieces from the membranes of the cell culture inserts were embedded into BEEM flat embedding molds (cat. #7004-01, Electron Microscopy Sciences, USA), so that they could be sectioned perpendicular to the membrane surface and thermally polymerized at 65 **°**C for 24 h. Ultra-thin sections with a thickness of 60 nm from polymerized resin blocks were sectioned using a Leica EM UC7 ultramicrotome (Leica Microsystems Inc. Canada), transferred on bare Cu grids, and post-stained with 2% uranyl acetate and Reinolds’ lead citrate for 10 minutes each. Sections were observed under a Hitachi H-7650 transmission electron microscope (Hitachi-High Technologies Canada) at 80 kV and imaged under a high definition electron multiplying charge coupled device (EMCCD) camera (XR111, Advanced Microscopy Techniques, USA).

### Immunofluorescence staining

For confocal imaging of claudin-1, tubulin and vimentin, cells were grown in FBS or in HS-containing media on poly-L-Lysine coated coverslips. Cells were washed with PBS, and fixed in 3.65% formaldehyde in PBS for 8 minutes at room temperature. After fixation cells were washed 3x with PBS, incubated with 0.2% Triton-X-100 for 2 minutes at room temperature, and washed again 3x with PBS, followed by a 2 hour blocking step in PBS containing 0.1% Tween-20 and 2% milk powder. Primary antibodies (anti-α-tubulin: Sigma-Aldrich, T6074; rabbit-anti-claudin-1: Invitrogen 71-7800; mouse anti-vimentin (v9): Abcam Ab8069), diluted in 0.1% Tween-20 PBS containing 2% milk, were added to the cells and incubated overnight at 4 °C. Cells were washed the next day, 3x, in PBS containing 0.1% Tween-20 followed by incubation with Alexa Fluor conjugated secondary antibodies AlexaFluor488 donkey anti-mouse IgG (Life Technologies, A21202), AlexaFluor594 donkey anti-mouse IgG (Life Technologies, A21203), AlexaFluor594 donkey anti-rabbit IgG (Life Technologies, A11012) for 45 minutes at room temperature. After washing the cells 3x in PBS 0.1% Tween-20 cells were incubated in a Hoechst stain, 1:3000 for 5 minutes and coverslips were mounted to slides using Fluoromount-G (SothernBiotech). Confocal images were obtained with a LSM 710 Axio Observer microscope (Carl Zeiss Inc.) using a 63x/1.40 NA Oil DIC Plan-Apochromat objective. Images were acquired as a z-stack series (with a distance of 0.24μm between slices) and are represented as a z-projection.

### Dextran diffusion studies

Cells were grown on collagen coated transwell dishes and in the case of HS-cultured cells allowed to differentiate, or in the case of FBS-cultured cells grown to confluency. Cells were then place in phenol red free media, and 1 μg/ml 70kDa dextran conjugated to Oregon green was added to the top compartment. Samples were taken every 30 minutes, for a total of 2 hours, from the bottom compartment to determine dextran diffusion rates. Orgeon green Fluorescence was measured on an Enspire 2300 Multilabel reader (Perkin Elmer) and compared to subconfluent FBS-cultured cells to determine maximal diffusion rates. Confluency of FBS-cultured cells was ensured by visualizing cells on filters under a conventional phase-contrast microscope, as well as measuring FBS diffusion rates on 3 consecutive days. The lowest diffusion rate of those 3 was used, if differences existed.

### NMR analysis of reporter metabolites

Target profiling and analysis of reporter metabolites by NMR was performed by Chenomx Inc, Edmonton, Canada. In short, FBS-cultured cells and confluent, differentiated HS-cultured cells were washed with DMEM and placed in DMEM (without serum) overnight. Supernatants of HS or FBS-cultured cells were collected, first filtered through 22μm filters to remove large debris. Internal standard solution (IS-1) was added to each sample, and the resulting mixture was vortexed for 30 seconds. Samples were then filtered through Nanosep 3K Omega microcentrifuge tubes to remove all proteins and other large complexes, and transferred to an NMR tube for data acquisition

NMR spectra were acquired on a Varian two-channel VNMRS 600 MHz NMR spectrometer equipped with an HX 5 mm probe. The pulse sequence used was a 1D-tnnoesy with a 990 ms presaturation on water and a 4 s acquisition time. Spectra were collected with 32 transients and 4 steady-state scans at 298 K. Spectra were tehn processed using the Processor module in Chenomx NMR Suite 8.0. Compounds were identified and quantified using the Profiler module in Chenomx NMR Suite 8.0 with the Chenomx Compound Library version 9, containing 332 compounds.

### RNA extraction and cDNA synthesis

Total RNA from cells was isolated using TRIzol reagent (Invitrogen) according to the manufacturer's instructions, and quantified by measuring the absorbance at 260 nm. RNA quality was determined by measuring the 260/280 ratio. Thereafter, first-strand cDNA synthesis was performed by using the High-Capacity cDNA reverse transcription kit (Applied Biosystems) according to the manufacturer’s instructions. Briefly, 1.5 μg of total RNA from each sample was added to a mix of 2.0 μl 10X RT buffer, 0.8 μl 25X dNTP mix (100 mM), 2.0 μl 10X RT random primers, 1.0 μl MultiScribe^TM^ reverse transcriptase, and 3.2 μl nuclease-free water. The final reaction mix was kept at 25°C for 10 min, heated to 37 °C for 120 min, heated for 85 °C for 5 sec, and finally cooled to 4 °C.

### Quantification by real time-PCR

Quantitative analysis of specific mRNA expression was performed by real time-PCR, by subjecting the resulting cDNA to PCR amplification using 96-well optical reaction plates in the Bio-rad CFX-96 Real-Time PCR System (Bio-rad). 25-μl reaction mix contained 0.1 μl of 10 μM forward primer and 0.1 μl of 10 μM reverse primer, 12.5 μl of SYBR Green Universal Mastermix, 11.05 μl of nuclease-free water, and 1.25 μl of cDNA sample. The primers used in the current study were chosen from previously published studies^40^. No-template controls were incorporated onto the same plate to test for the contamination of any assay reagents. An optical adhesive cover was used to seal the plate; thereafter, thermocycling conditions were initiated at 95°C for 10 min, followed by 40 PCR cycles of denaturation at 95°C for 15 sec, and annealing/extension at 60°C for 1 min. Dissociation curves were performed by the end of each cycle to confirm the specificity of the primers and the purity of the final PCR product.

### Real time-PCR Data analysis

The real time-PCR data were analyzed using the relative gene expression method. Briefly, the data are presented as the fold change in gene expression normalized to the endogenous reference gene (β-actin) and relative to the untreated control of the same time point.

### Measuring different P450s activities using cocktail substrates

The reaction was started by incubating live intact cells with 200 μL William’s E Medium containing a mixture of substrates including, 100 μM 7-methoxyresorufin (CYP1A2 substrate), 500 μM bupropion (CYP2B6 substrate), 20 μM paclitaxel (CYP2C8 substrate), 250 μM S-mephenytoin (CYP2C19 substrate), 15 μM dextromethorphan (CYP2D6 substrate), 200 μM testosterone (CYP3A substrate) in addition to 20 unites of sulfatase and 5000 units of β-glucuronidase. The reaction was stopped by the addition of 200 μL ice cold methanol containing 100 μM acetaminophen as internal standard for LC/MS analysis.

Metabolites formed were analyzed on a liquid chromatography – tandem mass spectrometry system, similar to a previously published method^41^. Selected reaction monitoring in the positive-ion electrospray ionization mode was performed for acetaminophen as an internal standard, resorufin (a metabolite of 7-methoxyresorufin produced by CYP1A2 activity), hydroxybupropion (a metabolite of bupropion produced by CYP2B6 activity), 6a-hydroxypacliataxel (a metabolite of paclitaxel produced by CYP2C8 activity), 4′-hydroxymephenytoin (a metabolite of S-mephenytoin produced by CYP2C19 activity), dextrorphan (a metabolite of dextromethorphan produced by CYP2D6 activity), and 6β-hydroxytestosterone (a metabolite of testosterone produced by CYP3A activity).

### Data availability, code availability

Microarray data were deposited in the GEO repository with Accession Number GSE87684 (GEO), other source data are available in the supplemental data. Code is available from the authors upon request.

### Statistical analysis

For calculation of significance of quantitative data, the Prism Statistics package was used. All experiments consisted of a minimum of 3 independent replicates (further detailed in figure legends), and statistical significance was calculated using ‘Student’s t-test (unpaired, two-tailed) for comparing two samples, or one-sided ANOVA/ Dunnett (with adjustments for multiple comparisons) for comparision of 3 or more samples. Values are depicted as means, with s.d. or s.e.m. as indicated in the legends. P-values of 0.05 or smaller were considered significant. P values ranges are depicted as asterisks: *: P< 0.05, **: P< 0.01, ***: P< 0.001, ****: P≤0.0001, n.s: P> 0.05

## Figure Legends

**Supplemental Data 1.**
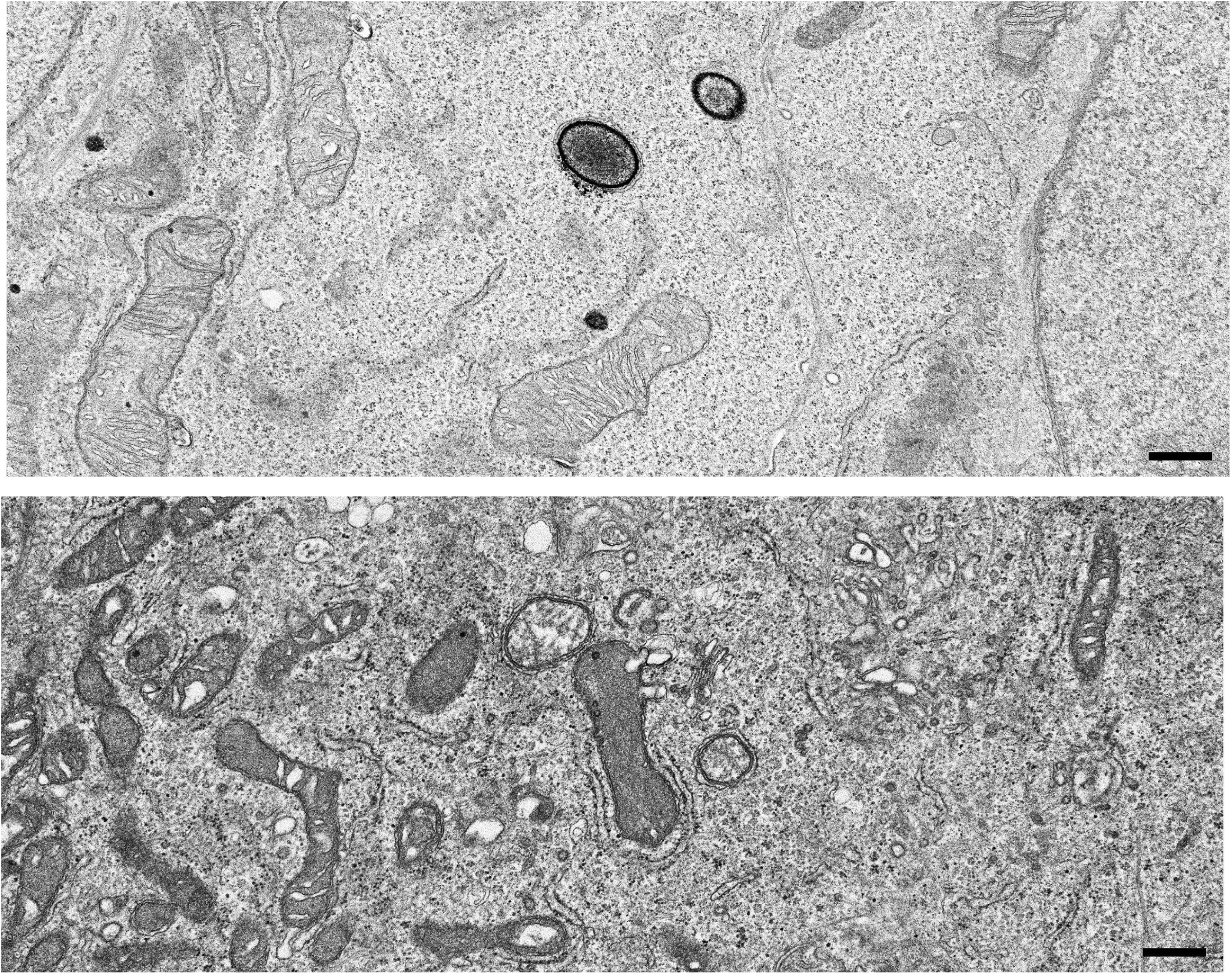
other morphological changes in HS cultured cells.

Additional morphological changes that were observed in the elelctron microscopic analysis include:

- the cytoplasm of HS cultured cells appears much more crowded than the cytoplasms of FBS cultured cells.
- mitochondiral morphology has drastically changed, as outlined in section 4 of the main document
- in general. organelles appeared more structured and organized, for example, the endop[lasmic reticulum shows a higher degree of organization, particularly around the mitochondira. We assume these regions represent the mitochondria associated membranes or MAM.
- An increase in vesicle transport and changes in the lysosomal pathways were also observed on the electron micrographs. Whereas in FBS cells early lysosomes (dark structures in the top figure) are abundant, but other components of the endosomal/lysosomal pathway were not detected, lysosomal structures were not seen on EM images of HS cultured cells. Instead, in HS cultured cells late endosomes, multivesicular bodies and autophagosome, potentially indicating that a more complete endosomal/ lysosomal pathway is operational in HS cultured cells.

**Supplemental Data 2.**
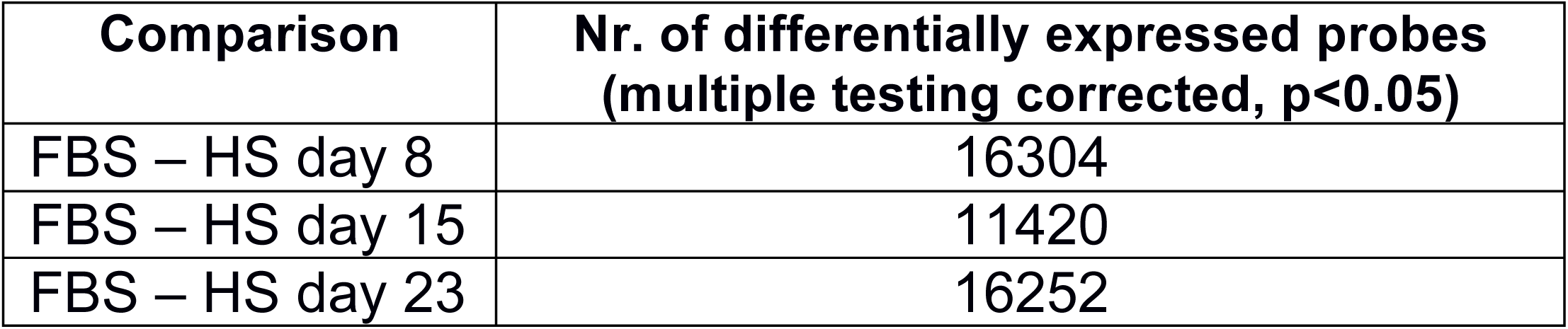
Transcriptional changes in HS-cultured cells compared to FBS-cultured cells.

**Supplemental data 3.**
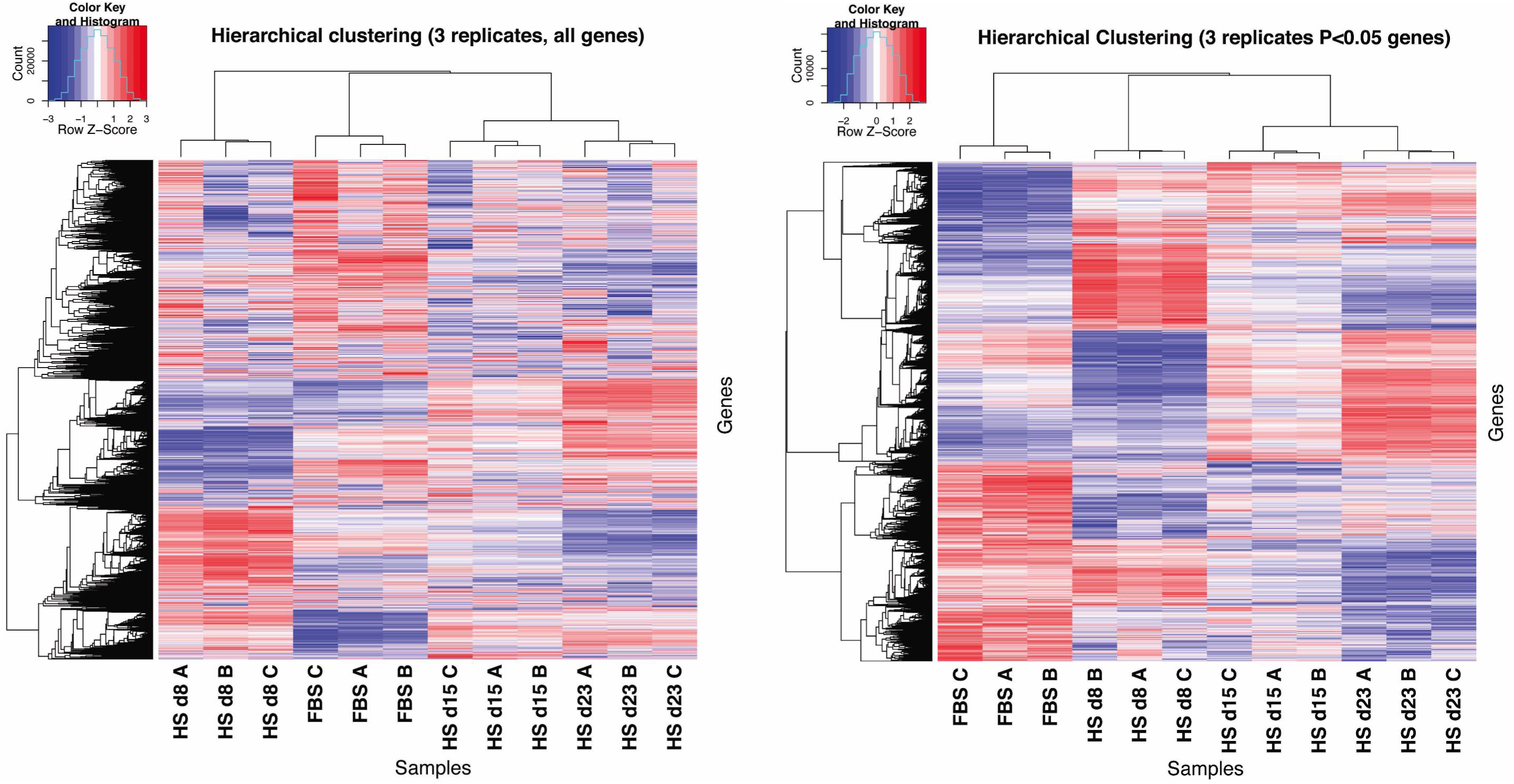
hierarchical clustering analysis.

Clustering is a machine learning algorithm that groups data that have a high degree of similarity together in a cluster. In this analysis, data were clustered, initially, by using hierarchical clustering as shown above, based on gene expression pattern similarity of the entire dataset (y-axis) as well as on patterns of expression over time (x-axis). The z-scores of the gene expression levels were used in order to focus on the expression variation pattern across the samples rather than on their absolute expression levels, which can vary strongly from gene to gene.

Hierarchical clustering of the entire data set is shown on the left and of transcripts that showed a significant change (p < 0.05) is shown on the right In both analyses the replicates clustered together (x-axis), as was also shown by the PCA analysis (figure 1), and 6 patterns emerged (y-axis) when the data with significant changes were considered (right panel). To further analyze the genes or processes that were associated with these 6 clusters, a variant of k-means clustering was used, PAM clustering (partitioning around medoids), to generate more clearly delineated clusters (see figure 3 in the main document).

**Supplemental data 5.**
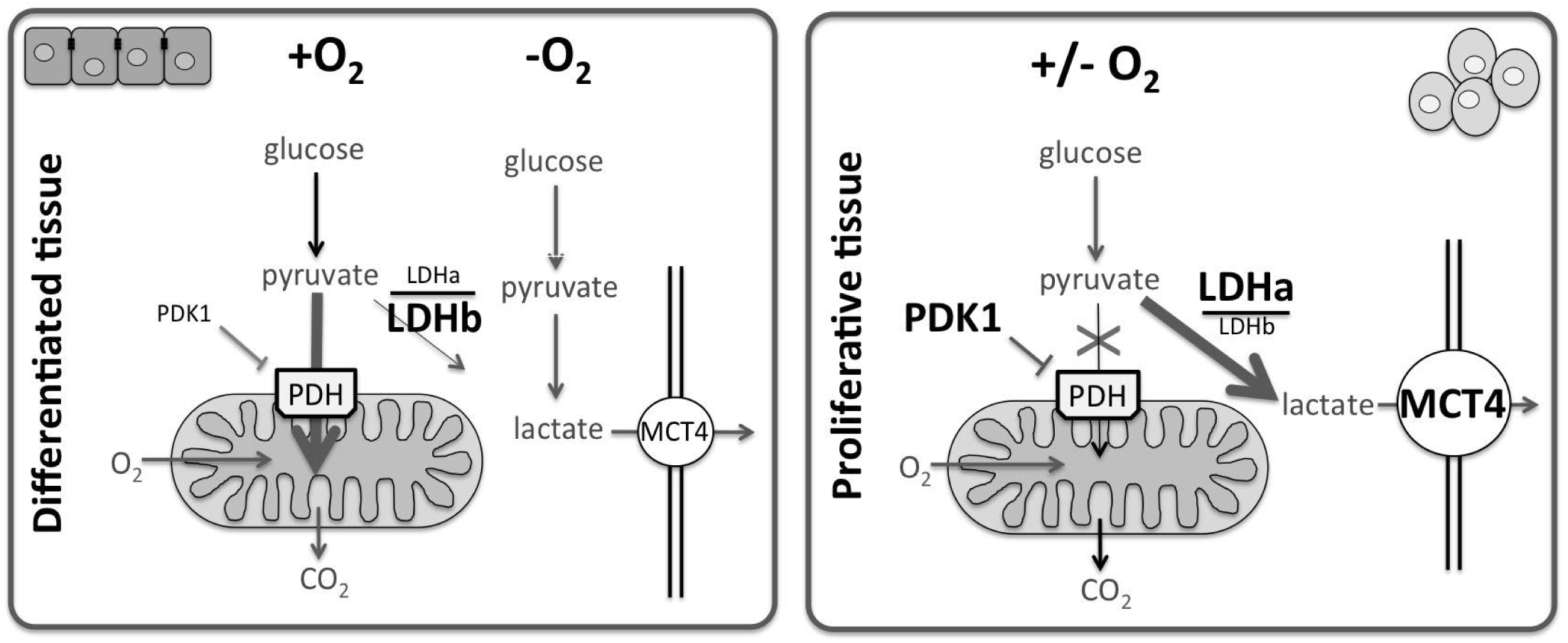
regulation of the Warburg effect.

Proliferating cells often display a metabolic profile that is referred to as ‘cancer metabolism’, first described by Otto Warburg in 1924. Cancer metabolism is typically characterized by reduced levels of oxidative phosphorylation and mitochondrial activity, higher dependence on aerobic glycolysis and glutaminolysis for ATP production, and increased generation of biosynthetic intermediates that are essential for the production of macromolecules (phospholipids, nucleotides, proteins) to support cell proliferation (reviewed in^3, 5-7^). The metabolic reprogamming that occurs during the Warburg effect is tightly regulated. The likely benefit for proliferative cells to adopt a Warburg-like metabolic profile, despite the much lower yield of ATP, is the conservation of pyruvate for the synthesis of lipids, nucleotides and amino acids, as building blocks for new cells ^1-4^.

Key regulators of the Warburg effect include:

* **PDK1** (pyruvate dehydrogenase kinase 1): in proliferative cells PDK1 is up-regulated, inhibiting PDH (pyruvate dehydrogenase), which transport pyruvate into the mitochondria, PDK1 upregualtion results in limiting the uptake of pyruvate into the mitochondria. *PDK1 is down-regulated in HS-cultured cells, in line with a non-proliferative character. PDH did not change.*

* **LDHA and LDHB:** lactate dehydrogenases A and B catalyze the conversion of pyruvate ot lactate (LDHA), vice versa (LDHB) High LDHA levels direct the conversion of pyruvate to lactate, whereas low LDHa levels, with higher LDHB levels favour the reverse reaction. *LDHA is decreased in HS cultured cells, whereas LDHB is increased, consistent with the ratio in differentiated tissue*

* **MCT4** (monocarboxylic acid transporter 4): MCT4 removes lactate from the cells, and is increased during aerobic glycolysis (right panel). *MCT4 is decreased in HS cultured cells, consistent with the reversal of the Warburg effect.*

**Supplemental Data 6:**
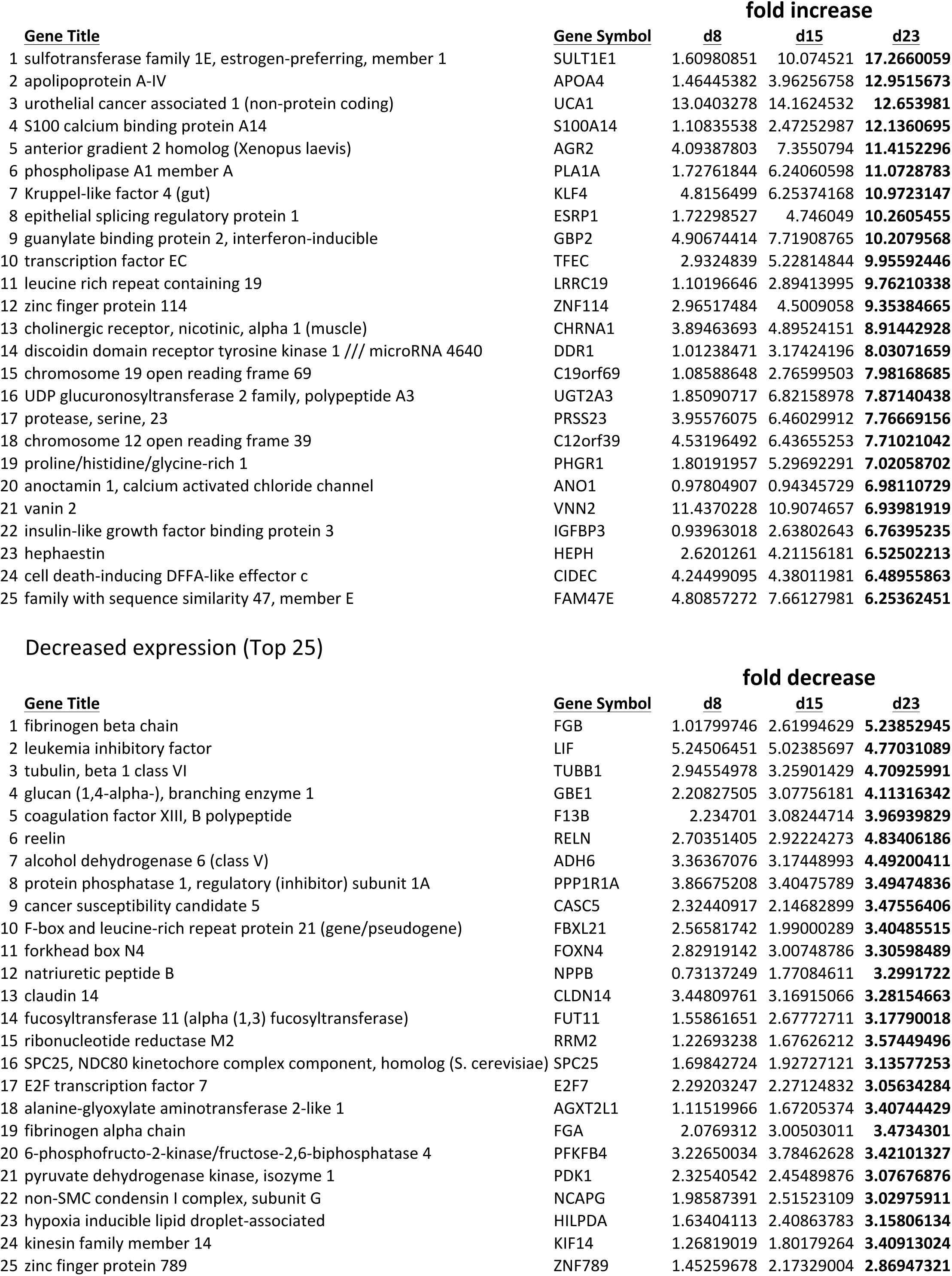
25 genes with highest increase or decrease in expression.

**Supplemental Data 7.**
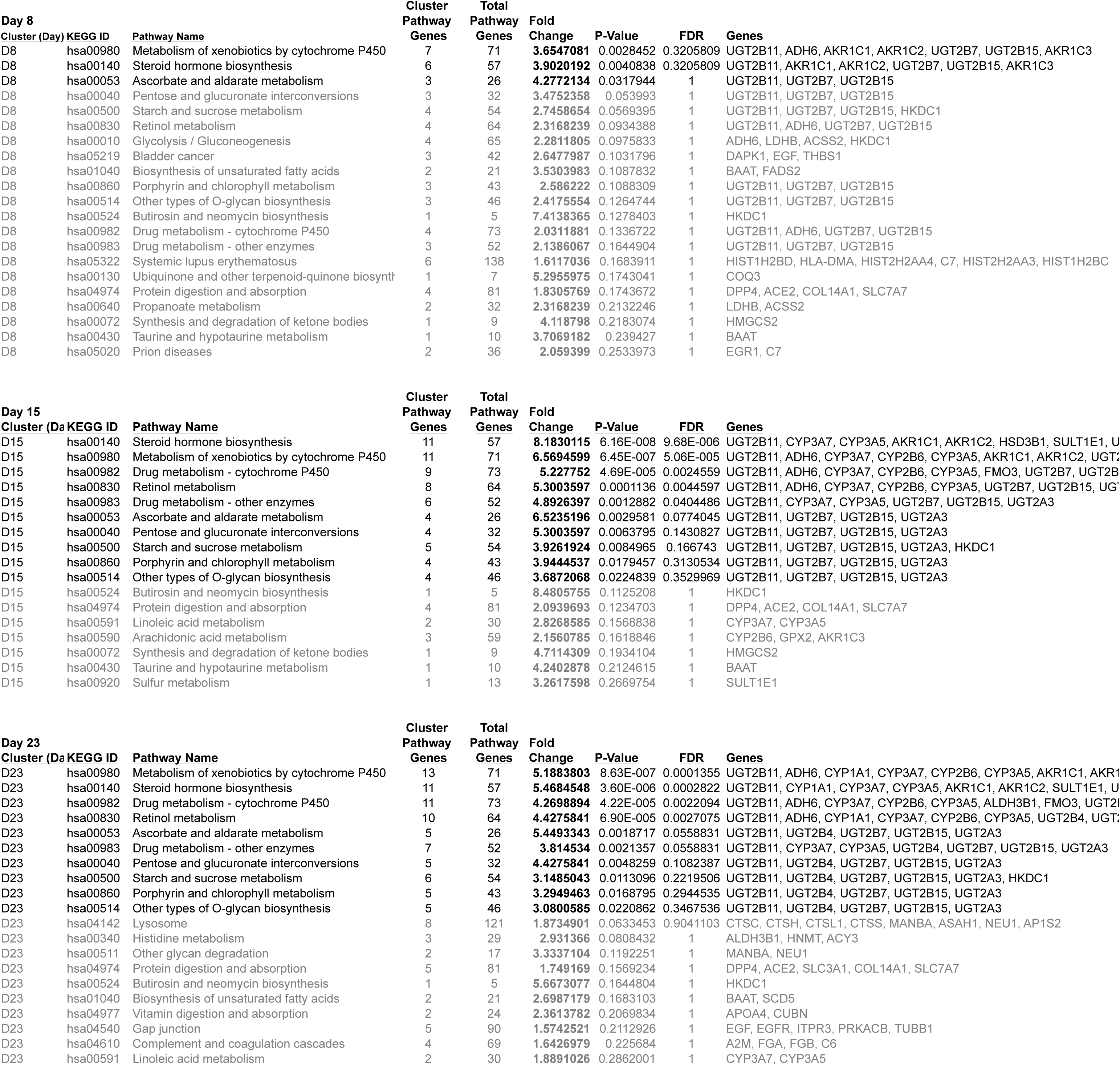
Differentially expressed KEGG pathways (p< 0.25) (referring to Table 1)

**Supplemental Data 8.**
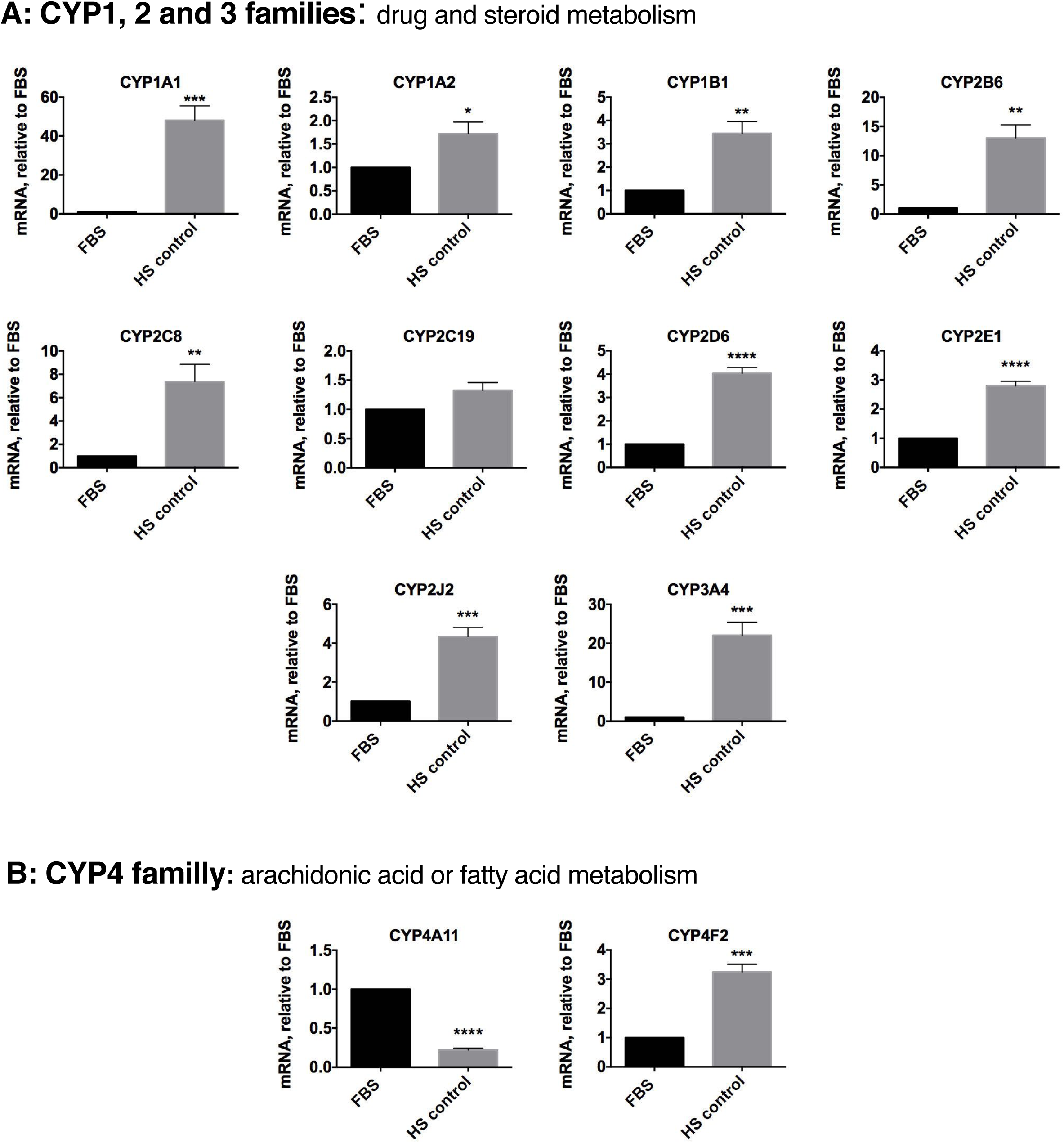
(referring to figure 6) mRNA levels determined by quantitative PCR of different cytochrome P450 genes.

**Supplemental Data 9.**
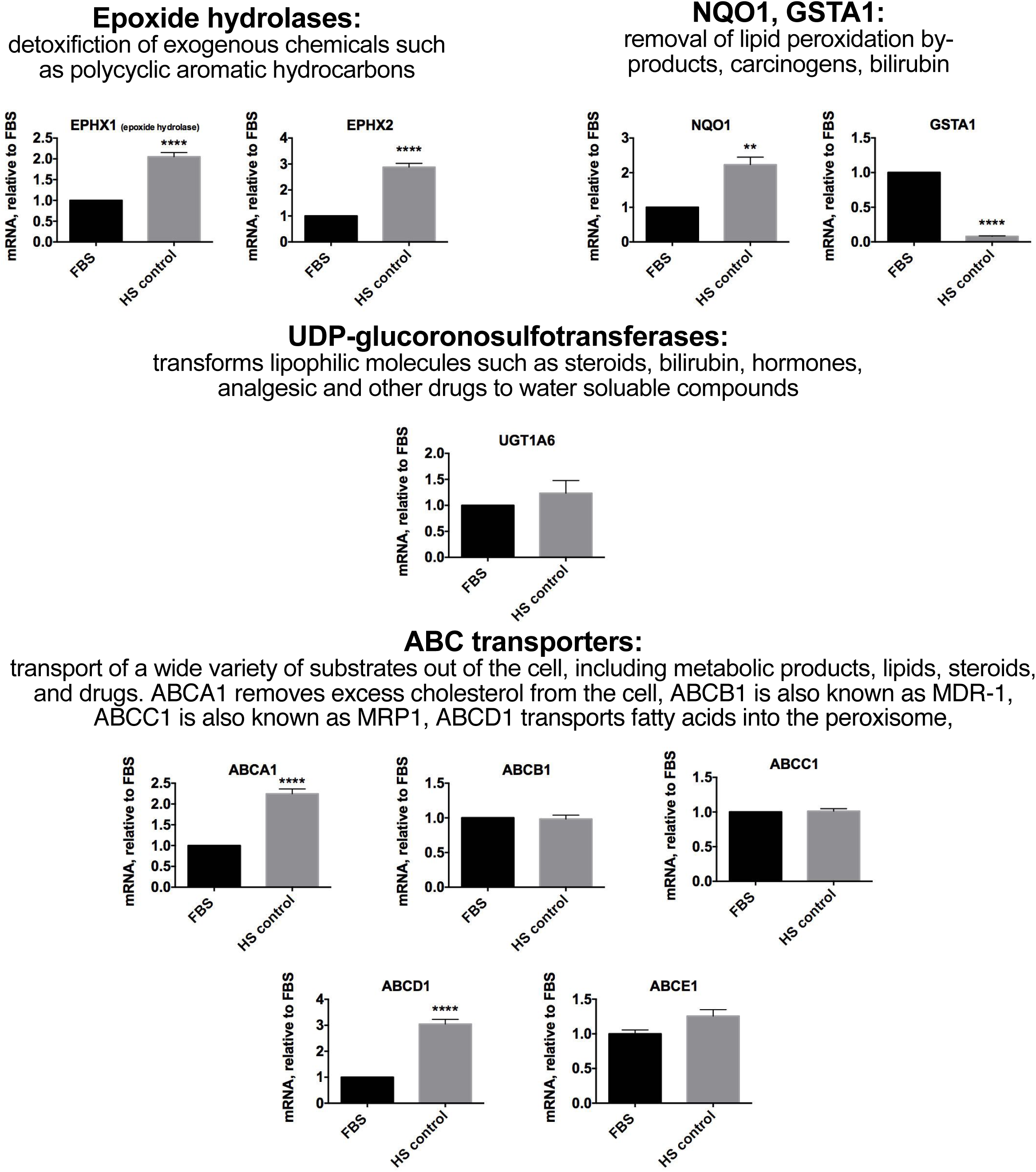
(referring to figure 6) mRNA levels determined by quantitative PCR of other factors involved in degradation or removal of xenobiotics

## References

1. Berg, J.M., Tymoczko, J.L. & Stryer, L. Section 30.2, Each Organ Has a Unique Metabolic Profile., in Biochemistry. (ed. W.H. Freeman) (MacMillan Learning, New York; 2002).

2. Van den Berghe, G. The role of the liver in metabolic homeostasis: implications for inborn errors of metabolism. Journal of Inherited Metabolic Diseases 14, 407’420 (1991).

3. Vander Heiden, M.G., Cantley, L.C. & Thompson, C.B. Understanding the Warburg effect: the metabolic requirements of cell proliferation. Science 324, 1029’1033 (2009).

4. Steenbergen, R.H. et al. Human serum leads to differentiation of human hepatoma cells, restoration of VLDL secretion and a 1000-fold increase in HCV JFH-1 titers. Hepatology (2013).

5. Cairns, R.A., Harris, I.S. & Mak, T.W. Regulation of cancer cell metabolism. Nat Rev Cancer 11, 85’95 (2011).

6. Lunt, S.Y. & Vander Heiden, M.G. Aerobic glycolysis: meeting the metabolic requirements of cell proliferation. Annu Rev Cell Dev Biol 27, 441’464 (2011).

7. Pavlova, N.N. & Thompson, C.B. The Emerging Hallmarks of Cancer Metabolism. Cell Metab 23, 27’47 (2016).

8. Hackenbrock, C.R. Chemical and physical fixation of isolated mitochondria in low-energy and high-energy states. Proc Natl Acad Sci U S A 61, 598’605 (1968).

9. Hackenbrock, C.R. States of activity and structure in mitochondrial membranes. Ann N Y Acad Sci 195, 492’505 (1972).

10. Hackenbrock, C.R. Energy-linked ultrastructural transformations in isolated liver mitochondria and mitoplasts. Preservation of configurations by freeze-cleaving compared to chemical fixation. J Cell Biol 53, 450’465 (1972).

11. Perkins, G.A. & Ellisman, M.H. Mitochondrial configurations in peripheral nerve suggest differential ATP production. J Struct Biol 173, 117’127 (2011).

12. Mannella, C.A. et al. Topology of the mitochondrial inner membrane: dynamics and bioenergetic implications. IUBMB Life 52, 93’100 (2001).

13. Rossignol, R. et al. Energy substrate modulates mitochondrial structure and oxidative capacity in cancer cells. Cancer research 64, 985’993 (2004).

14. Garcia-Perez, A.I. et al. Molecular crowding and viscosity as determinants of translational diffusion of metabolites in subcellular organelles. Arch Biochem Biophys 362, 329’338 (1999).

15. Lopez-Beltran, E.A., Mate, M.J. & Cerdan, S. Dynamics and environment of mitochondrial water as detected by 1H NMR. J Biol Chem 271, 10648’10653 (1996).

16. Sonnewald, U., Schousboe, A., Qu, H. & Waagepetersen, H.S. Intracellular metabolic compartmentation assessed by 13C magnetic resonance spectroscopy. Neurochem Int 45, 305’310 (2004).

17. Singaravelu, R. et al. Human serum activates CIDEB-mediated lipid droplet enlargement in hepatoma cells. Biochem Biophys Res Commun 441, 447’452 (2013).

18. Laffel, L. Ketone bodies: a review of physiology, pathophysiology and application of monitoring to diabetes. Diabetes Metab Res Rev 15, 412’426 (1999).

19. Pramfalk, C., Larsson, L., Härdfeldt, J., Eriksson, M. & Parini, P. Culturing of HepG2 cells with human serum improve their functionality and suitability in studies of lipid metabolism. Biochimica et Biophysica Acta (BBA) - Molecular and Cell Biology of Lipids 1861, 51’59 (2016).

20. Gey, G.O., Coffman, W.D. & Kubicek, M.T. Tissue culture studies of the proliferative capacity of cervical carcinoma and normal epithelium. . Cancer research 12, 264’265 (1952).

21. Frezza, C. & Gottlieb, E. Mitochondria in cancer: not just innocent bystanders. Semin Cancer Biol 19, 4’11 (2009).

22. Matoba, S. et al. p53 regulates mitochondrial respiration. Science 312, 1650’1653 (2006).

23. Bressac, B. et al. Abnormal structure and expression of p53 gene in human hepatocellular carcinoma. Proc Natl Acad Sci U S A 87, 1973’1977 (1990).

24. Puisieux, A. et al. Retinoblastoma and p53 tumor suppressor genes in human hepatoma cell lines. FASEB J 7, 1407’1413 (1993).

25. Jiao, X.-J., Steenbergen, R.H.G. & Tyrrell, D.L. The use of human umbilical cord blood serum is beneficial for the continuous production of hepatitis C virus. Journal of General Virology (epub ahead of print) (2016).

26. Ang, L.P. et al. Ex vivo expansion of conjunctival and limbal epithelial cells using cord blood serum-supplemented culture medium. Invest Ophthalmol Vis Sci 52, 6138’6147 (2011).

27. Ayatollahi, M., Soleimani, M., Geramizadeh, B. & Imanieh, M.H. Insulin-like growth factor 1 (IGF-I) improves hepatic differentiation of human bone marrow-derived mesenchymal stem cells. Cell biology international 35, 1169’1176 (2011).

28. Magner, N.L. et al. Insulin and IGFs enhance hepatocyte differentiation from human embryonic stem cells via the PI3K/AKT pathway. Stem Cells 31, 2095’2103 (2013).

29. Regel, I. et al. IGFBP3 impedes aggressive growth of pediatric liver cancer and is epigenetically silenced in vascular invasive and metastatic tumors. Mol Cancer 11, 9 (2012).

30. Stead, D. & Welch, V.A. Lipid composition of bovine serum lipoproteins. J Dairy Sci 58, 122’127 (1975).

31. Dashti, N., Smith, E.A. & Alaupovic, P. Increased production of apolipoprotein B and its lipoproteins by oleic acid in Caco-2 cells. Journal of lipid research 31, 113’123 (1990).

32. Ihaka, R. & Gentleman, R.R : A language for data analysis and graphics. Journal of Computational and Graphical Statistics 5, 299’314 (1996).

33. Gentleman, R.C. et al. Bioconductor: open software development for computational biology and bioinformatics. Genome biology 5, 1’16 (2004).

34. Irizarry, R.A. et al. Exploration, normalization, and summaries of high density oligonucleotide array probe level data. Biostatistics 4, 249’264 (2003).

35. Ritchie, M.E. et al. limma powers differential expression analyses for RNA-sequencing and microarray studies. Nucleic Acids Research 43, e47 (2015).

36. Young, M.D., Wakefield, M.J., Smyth, G.K. & Oshlack, A. Gene ontology analysis for RNA-seq: accounting for selection bias. Genome biology 11, 1’12 (2010).

37. Thiele, I. et al. A community-driven global reconstruction of human metabolism. Nature biotechnology 31, 419’425 (2013).

38. Jensen, P.A. & Papin, J.A. Functional integration of a metabolic network model and expression data without arbitrary thresholding. Bioinformatics 27, 541’547 (2011).

39. Romero, P. et al. Computational prediction of human metabolic pathways from the complete human genome. Genome biology 6, R2 (2005).

40. Nishimura, M., Yoshitsugu, H., Naito, S. & Hiraoka, I. Evaluation of gene induction of drug-metabolizing enzymes and transporters in primary culture of human hepatocytes using high-sensitivity real-time reverse transcription PCR. Yakugaku Zasshi 122, 339’361 (2002).

41. Pillai, V.C., Strom, S.C., Caritis, S.N. & Venkataramanan, R. A sensitive and specific CYP cocktail assay for the simultaneous assessment of human cytochrome P450 activities in primary cultures of human hepatocytes using LCMS/MS. J Pharm Biomed Anal 74, 126’132 (2013).

## References

1. Cairns, R.A., Harris, I.S. & Mak, T.W. Regulation of cancer cell metabolism. Nat Rev Cancer 11, 85-95 (2011).

2. Lunt, S.Y. & Vander Heiden, M.G. Aerobic glycolysis: meeting the metabolic requirements of cell proliferation. Annu Rev Cell Dev Biol 27, 441-464 (2011).

3. Pavlova, N.N. & Thompson, C.B. The Emerging Hallmarks of Cancer Metabolism. Cell Metab 23, 27-47 (2016).

4. Vander Heiden, M.G., Cantley, L.C. & Thompson, C.B. Understanding the Warburg effect: the metabolic requirements of cell proliferation. Science 324, 1029-1033 (2009).

